# ATF4-dependent upregulation of Bruno 1 remodels P-bodies to selectively protect mRNAs during ER stress throughout *Drosophila melanogaster* oogenesis

**DOI:** 10.64898/2026.04.01.715972

**Authors:** Samantha N. Milano, Livia V. Bayer, Julie J. Ko, Gwendolyn S. Posner, Albina H. Granovsky, Diana P. Bratu

## Abstract

P-bodies are cytoplasmic membraneless organelles involved in mRNA storage, yet their role in cellular stress responses remains poorly understood. Here, we demonstrate that P-bodies are rapidly and selectively remodeled during the early response to endoplasmic reticulum (ER) stress in *D. melanogaster* oogenesis, positioning them as key early stress responders. Notably, this remodeling occurs within minutes of stress induction and precedes stress granule formation. This early remodeling is characterized by changes in P-body morphology and internal organization and promotes selective mRNA regulation. Specifically, ER stress leads to the recruitment and stabilization of maternal mRNAs and those encoding P-body components, while transcripts not associated with P-bodies are degraded. These observations indicate that P-body remodeling is not merely structural but functionally linked to the selective preservation of mRNA populations during stress. Mechanistically, we find that this process is driven by transcriptional upregulation of the RNA-binding protein, Bruno 1, downstream of ATF4-dependent stress signaling, thereby establishing a direct connection between the unfolded protein response and condensate regulation. Consistent with this model, loss of Bruno 1 abolishes, whereas its overexpression enhances P-body remodeling, demonstrating that stress-induced changes in RNA binding protein levels can actively reprogram condensate properties. Together, our findings reveal that P-bodies function as dynamic, stress-responsive hubs that integrate transcriptional signaling with post-transcriptional control, enabling the selective preservation of essential mRNAs during ER stress. More broadly, this work uncovers a previously unrecognized mechanism by which stress signaling pathways reorganize cytoplasmic architecture to shape mRNA fate.

## Introduction

Cells rely on tightly coordinated mechanisms to modulate mRNA fate during cellular stress, such as heat shock, nutrient deprivation, or hypoxia (Fulda et al. 2010). To survive these perturbations, cells must rapidly adjust mRNA stability, localization, and translation (Srikantan and Gorospe 2010; Yamasaki and Anderson 2008). In many cases, translationally stalled mRNAs are sequestered within well-characterized cytoplasmic membraneless organelles (MLOs) called stress granules, where they are preserved until conditions improve and translation can resume (Protter and Parker 2016). While the role of stress granules in stress adaptation has been extensively studied, the contribution of P-bodies to stress, more specifically, to the early response to endoplasmic reticulum (ER) stress remains poorly understood.

Accumulation of unfolded or misfolded proteins within the ER lumen causes ER stress and activates the unfolded protein response (UPR) (Read and Schröder 2021). The UPR engages three primary signaling branches mediated by XBP1 (X-box binding protein 1), ATF4 and ATF6 (activating transcription factors 4 and 6), which together act as transcription factors and reprogram gene expression to increase ER chaperone production, reduce ER load with general mRNA degradation, suppress global protein synthesis, and restore ER proteostasis (Hetz et al. 2020; Cao 2012). The ER stress response is generally cytoprotective; however, prolonged or unresolved ER stress can become detrimental, leading to autophagy or chronic pathological states (Yoshida 2007; Yorimitsu and Klionsky 2007). As such, persistent ER stress is implicated in the etiology of numerous diseases, including Alzheimer’s disease, Parkinson’s disease, cancer, and diabetes (Ajoolabady et al. 2022; Mou et al. 2020; Yadav et al. 2014; Cnop et al. 2012). Understanding how cells coordinate the ER stress response at the level of mRNA regulation is therefore of fundamental biological and clinical importance.

Prior work from our group and others indicates a functional connection between the ER and P-bodies (Milano et al. 2024; Jason E. Lee et al. 2020; Kilchert et al. 2010). P-bodies are conserved across species and cell types and are constitutively present in the cytoplasm, positioning them to act as early responders to ER stress (Luo et al. 2018). These MLOs primarily house translationally repressed transcripts and function as organizational hubs in which multiple mRNA species can be recruited, stored, and released in a regulated manner (Standart and Weil 2018). P-bodies are highly dynamic structures that can alter their mRNA content in response to cellular cues, thereby promoting selective transcript storage or release depending on the cellular environment (Sankaranarayanan et al. 2021).

P-bodies form via liquid-liquid phase separation (LLPS), which occurs when local concentration of mRNAs and associated proteins become sufficiently enriched to favor the formation of a distinct condensed phase (Banani et al. 2017; Brangwynne et al. 2009; Elbaum-Garfinkle et al. 2015). The molecular composition of these condensates influences their emergent material properties, which are governed by networks of weak, multivalent interactions, including protein:protein, RNA:RNA, and RNA:protein interactions (Mittag and Parker 2018; Borcherds et al. 2021; Wadsworth et al. 2024).

P-bodies exist along a continuum of material states ranging from liquid-like to more solid-like. *In vitro*, nascent condensates often exhibit liquid-like properties but can progressively mature into more rigid assemblies over time as intracondensate interactions stabilize (Jawerth et al. 2020). *In vivo*, cells appear to actively regulate these material properties, preventing aberrant solidification and maintaining functional condensate dynamics (Y. Lin et al. 2015; Jain et al. 2016; Kroschwald et al. 2015). Emerging evidence suggests that P-body material state influences function: more solid-like P-bodies may preferentially support long-term transcript storage, whereas more liquid-like condensates may facilitate rapid transcript exchange, recruitment, and release for translation or degradation (Sankaranarayanan et al. 2021; Milano et al. 2025).

P-body material properties are partially determined by the composition of their constituent proteins. Many P-body proteins contain intrinsically disordered regions (IDRs), which promote LLPS (Schütz et al. 2017; Protter et al. 2018). In *D. melanogaster*, several conserved P-body associated proteins have been identified, including: Me31B (DDX6), Trailer Hitch (Lsm14 homolog) Cup (eIF4E-binding protein), and Bruno 1 (a member of the CELF family). Me31B is a DEAD-box RNA helicase that represses translation and exhibits high interaction valency, enabling multivalent protein:protein and protein:RNA interactions (De Valoir et al. 1991; Nakamura et al. 2001). Trailer Hitch (Tral) is an RNA-binding protein enriched IDRs, which promote phase separation and modulate the emergent material properties of biomolecular condensates (Tritschler et al. 2008; Sankaranarayanan et al. 2021). Cup is a large IDR-containing protein that binds eIF4E and competitively inhibits assembly of the translation initiation complex, thereby preventing ribosome recruitment. Cup is recruited to target mRNAs independently as well as via Bruno 1, which recognizes Bruno response elements (BREs) within transcripts (Bayer et al. 2023; Piccioni et al. 2005; Kim-Ha et al. 1995). Collectively, these proteins, together with associated mRNAs, drive the formation of phase-separated P-bodies that function as selective hubs for post-transcriptional gene regulation.

Here, we investigate the previously undefined role of P-bodies in the cellular response to ER stress. The contribution of post-transcriptional regulation to maintaining cellular homeostasis during the earliest phases of ER stress remains poorly understood. Thus, studying the early stages of stress response, when timely changes in mRNA fate can shape adaptive or pathological outcomes is of value. Interestingly, we find that P-bodies respond rapidly to ER stress, prior to stress granule formation, by undergoing changes in material state and selectively accumulating translationally repressed mRNAs. This response promotes the stabilization of specific mRNAs and depends on signaling through the ATF4 branch of the UPR and upregulation of Bruno 1. Together, our findings identify P-bodies as early regulators of mRNA fate during ER stress and reveal a previously unappreciated layer of post-transcriptional control in the UPR.

## Results

### P-body morphology rapidly changes upon ER stress induction

To investigate how ER stress influences P-body organization, we utilized *D. melanogaster*, a well-established system for dissecting ER stress responses due to the high conservation of key regulatory pathways (Ryoo and Steller 2007; Ryoo 2015). We focused on egg chambers, as the female germline has long served as a powerful model for studying RNA regulation and biomolecular condensates. Each egg chamber comprises 16 interconnected germline cells encapsulated by a layer of somatic follicle cells. Among these germline cells, one differentiates into a transcriptionally silent oocyte, while the remaining 15 become transcriptionally active nurse cells that synthesize and supply all maternal mRNAs required for oocyte development (Fig. 1A) (McLaughlin and Bratu 2015). The high transcriptional output of nurse cells, coupled with the need for extensive post-transcriptional regulation and long-range mRNA transport to the oocyte, makes this system particularly well suited for studying P-body dynamics. Development initiates in the germarium and proceeds through 14 morphologically defined stages prior to embryogenesis. These stages are broadly categorized as early (stages 1-5), mid (stages 6-8), and late (stages 9-14) oogenesis. For consistency, most of our quantitative analyses were performed in nurse cells from mid-stage egg chambers (Fig. 1A, black box).

**Figure 1:**
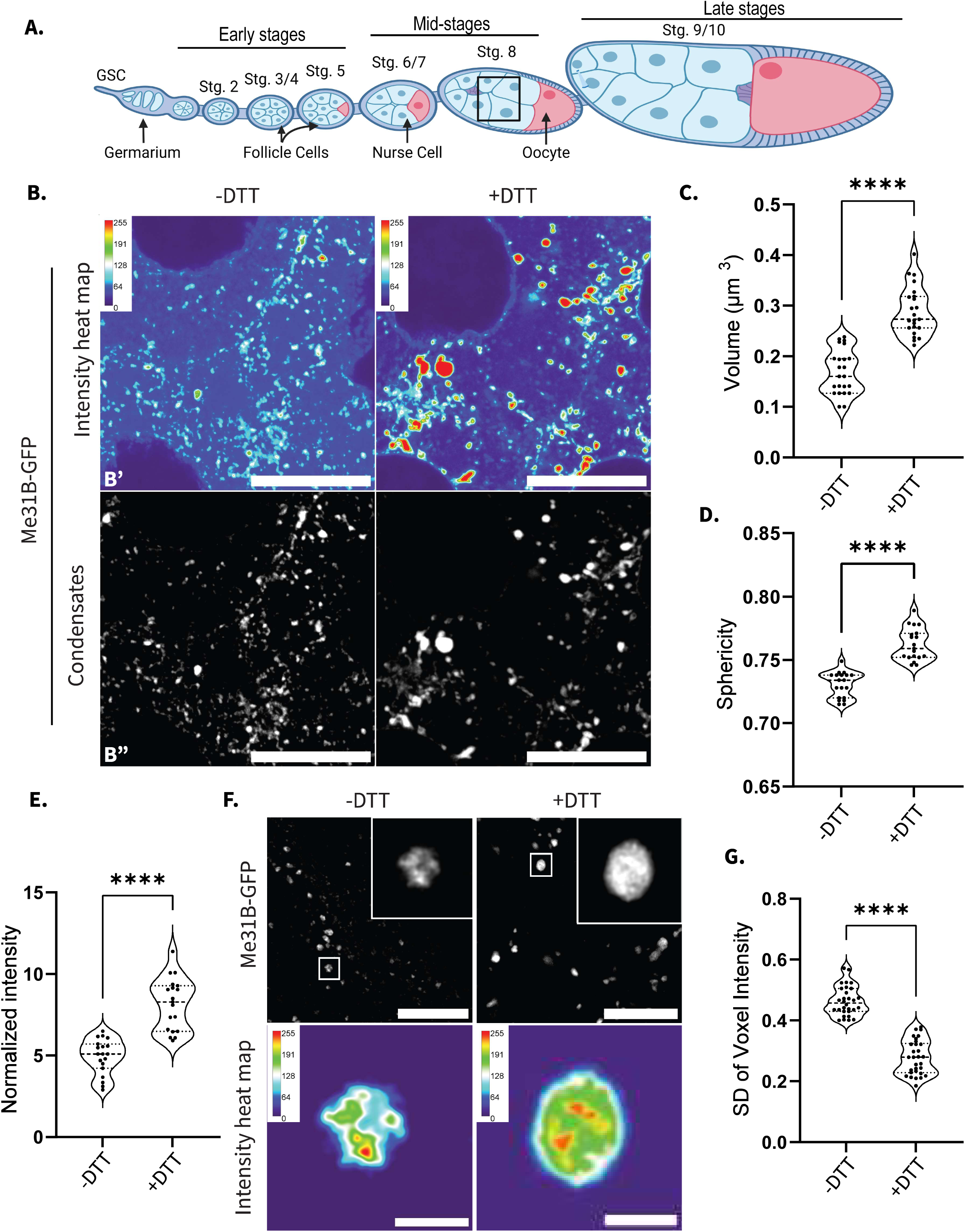
P-body morphology rapidly changes upon ER stress induction. **(A)** Schematic of a fly ovariole. Developmental stages progress from left to right. The oocyte (pink) and the nurse cells (light blue) are surrounded by somatic follicles (darker blue). The outlined ROI (black square) indicates the nurse cell region visualized for all zoomed image panels in this study. **(B)** Endogenously expressed Me31B-GFP visualized **(B’)** with an intensity heat map and **(B”)** with the cytoplasmic signal removed. Images are XY projections of 5 optical Z slices of 0.3µm. Scale bars are 20µm. **(C)** Volume quantifications comparing Me31B-GFP condensates in −DTT and +DTT incubated egg chambers (n =23). **(D)** Sphericity quantifications comparing Me31B-GFP condensates in −DTT and +DTT incubated egg chambers (n =19). **(E)** Voxel intensity quantifications comparing Me31B-GFP condensates in −DTT and +DTT incubated egg chambers (n =19). **(F)** Visualization of endogenous Me31B-GFP via STED. Images are XY projections of 5 optical Z slices of 0.22µm. Scale bar is 5µm (1µm for the zoomed inset). **(G)** Standard deviation of voxel intensity values for Me31B-GFP condensates in −DTT and +DTT incubated egg chambers (n=30). For all plots, each data point represents the average value of all P-bodies detected in an image. Significance was assessed using Mann-Whitney statistical tests. Error bars represent standard deviation. **** P < .0001.

To induce ER stress, ovaries were dissected and incubated *ex vivo* in Schneider’s media supplemented with 5 mM dithiothreitol (DTT), a well-established ER stressor that disrupts disulfide bond formation and activates the unfolded protein response (UPR) (Schröder 2008). P-bodies were visualized with endogenously tagged Me31B-GFP. Although Me31B is highly enriched within P-bodies, it is also distributed throughout the cytoplasm; notably, studies of its mammalian homolog have shown that more than 80% of the protein resides outside of condensates (Ernoult-Lange et al. 2012). To enhance visualization of the P-body associated signal, we removed the diffuse cytoplasmic Me31B signal (voxel intensities below 70) and displayed only condensate-localized Me31B throughout the study (Fig. 1B).

Following 30 minutes of DTT treatment (+DTT), Me31B-marked condensates exhibited a ∼71% increase in volume (from 0.169 µm³ to 0.289 µm³) and a ∼4.36% increase in sphericity (from 0.730 to 0.762) relative to control egg chambers incubated in Schneider’s media alone (−DTT) (Fig. 1C, D). Even modest increases in sphericity have been linked to reduced intracondensate bonds and are consistent with remodeling of condensate architecture (Sankaranarayanan et al. 2021; Bayer et al. 2023).

To confirm that these effects were due to ER stress rather than a DTT-specific artifact, we induced ER stress using an independent mechanism. Treatment with 1 μM Thapsigargin, which perturbs ER calcium homeostasis, similarly resulted in a ∼94% increase in P-body volume (from 0.168µm^3^ to 0.326µm^3^) and a ∼8.90% increase in sphericity (from 0.730 to 0.795) after 30 minutes (Fig. S1A-C) (Inesi and Sagara 1992).

We next asked whether the observed increase in P-body size reflected elevated Me31B protein levels or a redistribution of existing protein. Western blot analysis revealed that total Me31B protein levels were unchanged following ER stress induction, indicating that the morphological changes occur independently of altered Me31B expression levels (Fig. S1D). Consistent with the observed increase in condensate size, normalized Me31B-GFP intensity within P-bodies also increased by ∼62.29% following ER stress, further suggesting enhanced partitioning of Me31B into condensates rather than increased protein abundance (Fig. 1E).

To further confirm that these effects reflected general P-body remodeling rather than protein-specific behavior, we examined additional P-body components, Cup and Tral. Endogenously expressed Cup-YFP labeled condensates increased in volume by ∼68.85% and in sphericity by ∼4.74% upon ER stress induction (Fig. S1E-G). Similarly, endogenously expressed Tral-RFP condensates increased in volume by ∼128.09% and in sphericity by ∼8.59%, supporting the conclusion that ER stress broadly alters P-body morphology (Fig. S1H-J).

To investigate whether these morphological changes were accompanied by alterations in internal P-body organization, we employed STED super-resolution microscopy to resolve intracondensate structure (Fig. 1F). ER stress induction led to a marked change in the spatial distribution of Me31B-GFP fluorescent signal within individual P-bodies. To quantify this, we measured the standard deviation (SD) of voxel intensities within each condensate. Highly structured condensates exhibit heterogeneous fluorescence intensities and therefore higher voxel intensity SDs, whereas less internally structured condensates have fewer fluorescence peaks and valleys and thus exhibit lower internal voxel intensity SDs (Shiina 2019; Milano et al. 2024). Using this metric, we found that ER stress reduced the SD of voxel intensity within Me31B-GFP marked P-bodies by ∼39.96%, indicating a smoother and more homogeneous internal organization (Fig. 1G). Together this data suggests that P-bodies rapidly remodel their morphology and intracondensate organization within 30 minutes of ER stress onset.

### P-body remodeling during ER stress precedes stress granule assembly

Because P-bodies share multiple components and biophysical properties with stress granules, we next asked whether the observed changes in P-body morphology were secondary to stress granule formation. To address this, we examined egg chambers expressing endogenously tagged Rasputin (Rin)-GFP and performed immunostaining for eIF4G, both *bona fide* stress granule markers (Fig. 2A) (Buddika et al. 2020). Notably, we did not detect condensation of either Rin-GFP or eIF4G following 30 minutes of DTT treatment.

**Figure 2:**
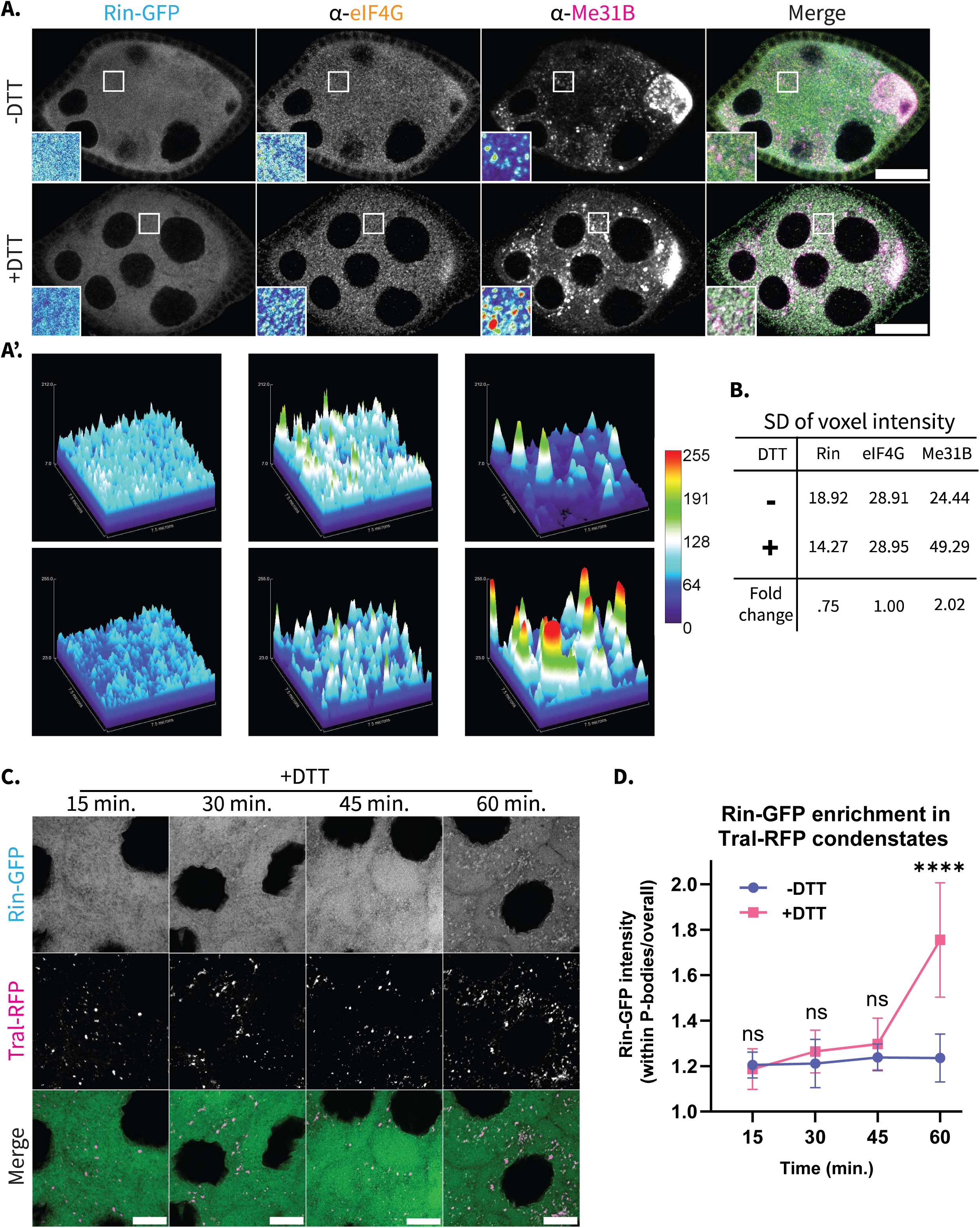
P-body remodeling during ER stress precedes stress granule assembly. **(A)** Endogenously expressed Rin-GFP covisualized with immunolabeled eIF4G and Me31B in egg chambers incubated *ex vivo* in Schneider’s media alone (-DTT) or in Schneider’s media supplemented with DTT (+DTT). Images are XY projections of 5 optical Z slices of 0.3µm. Scale bars are 50µm. **(A’)** 3D intensity maps of selected ROIs in **(A)**. **(B)** SD of voxel intensity of ROIs in **(A’)**. **(C)** Covisualization of endogenously expressed Rin-GFP and Tral-RFP in +DTT incubated egg chambers over the course of 60 min. Images are XY projections of 5 optical Z slices of 0.3µm. Scale bars are 20µm. **(D)** Quantification of Rin-GFP enrichment within Tral-RFP labeled P-bodies in −DTT and +DTT incubated egg chambers over the course of 60 min (n =20). For all plots, each data point represents the average value of all P-bodies detected in an image. Significance was assessed using Mann-Whitney statistical tests. Error bars represent standard deviation. **** P < .0001.

To quantitatively assess potential changes in protein distribution, we measured the SD of voxel intensity within defined regions of interest (ROIs), enabling comparison of spatial heterogeneity across conditions (Fig. 2A’). Consistent with our qualitative observations, eIF4G signal distribution remained unchanged between control and DTT-treated egg chambers. Similarly, Rin-GFP signal SD exhibited a modest decrease to 0.75-fold relative to control. In contrast, Me31B signal SD increased by ∼2.02-fold under DTT treatment, consistent with the pronounced morphological changes observed in P-bodies (Fig. 2B). Together, this data indicate that ER stress induces early remodeling of P-bodies in the absence of detectable stress granule assembly.

We next sought to define the temporal relationship between P-body remodeling and stress granule formation following ER stress. Previous studies have suggested that stress granules can nucleate from pre-existing P-bodies (Buchan et al. 2008). To test whether this occurs under ER stress conditions, we performed a time-course analysis co-visualizing Tral-RFP and Rin-GFP (Fig. 2C, S2A). Rin-GFP did not exhibit increased recruitment to Tral-positive condensates at early time points, and enrichment was only observed after 60 minutes of ER stress induction, at which point Rin-GFP association increased by ∼42.01% (Fig. 2D). Collectively, these results demonstrate that P-bodies undergo rapid morphological and organizational remodeling during the early phase of ER stress, preceding the detectable assembly of stress granules.

### ER stress leads to changes in P-body composition

Previous studies have indicated that changes in P-body intracondensate organization can lead to a change in P-body composition (Sankaranarayanan et al. 2021; Milano et al. 2025). As we observed clear changes in P-body organization within 30 minutes of ER stress, we next asked if the mRNA composition of P-bodies also shifted within this timeframe. We chose to explore three distinct classes of mRNAs in our study: (1) maternal mRNAs which normally localize to P-bodies (*oskar, bicoid,* and *nanos)*, (2) the mRNAs of P-body protein components (*me31B, cup,* and *bruno1*), and (3) non-P-body mRNAs (*armi* and *glorund*). To determine the localization of each mRNAs in our tissue, we created a grouped comparison of the three classes by generating a series of Vantage plots using Imaris software (Fig. 3A). These plots allow for the quantification of mRNA partitioning into P-bodies (puncta with the shortest distances to P-bodies of 0 µm or less localize inside while shortest distances larger than 0 µm localize outside P-bodies). Maternal mRNAs showed a much smaller average shortest distance to P-bodies than either other class. This was expected as these mRNAs are known to localize to P-bodies while the other two classes of mRNAs have not been shown to display this type of localization. Next, we plotted mRNA puncta volume against the shortest distance to Me31B-GFP marked P-bodies (Fig. S3A-H). Notably, mRNA particles with shorter distances to P-bodies displayed substantially larger volumes than particles with longer shortest distances to P-bodies, indicating the accumulation of multiple mRNA molecules within these condensates.

**Figure 3:**
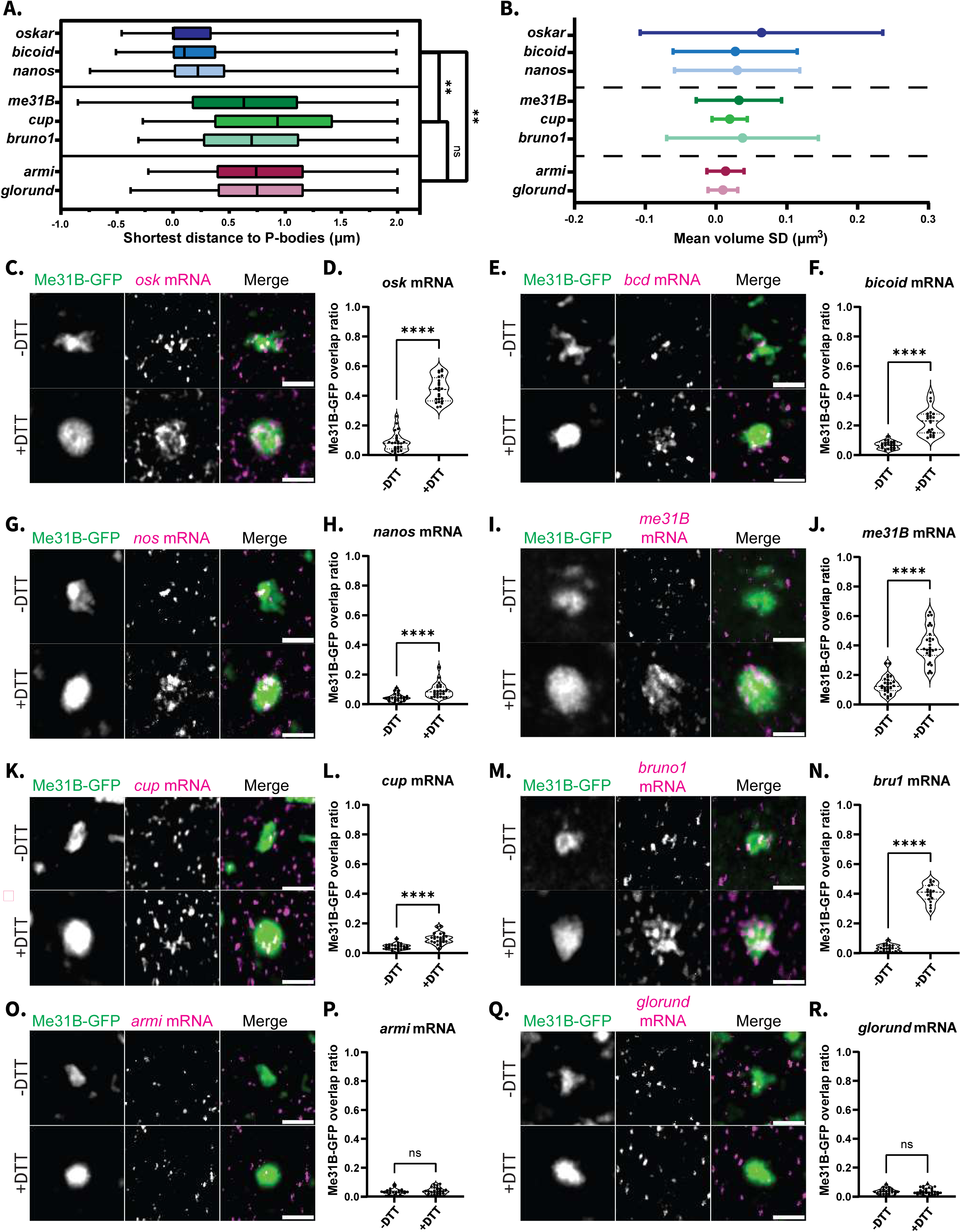
ER stress leads to changes in P-body composition. **(A)** Comparisons between the shortest distance to P-body calculations for three distinct classes of mRNA (shades of blue = maternal mRNAs; shades of green = P-body component mRNAs; shades of pink = non-P-body associated mRNAs) (n= 3 images, all mRNA particles are represented from each image). Significance was assessed with Nested one-way ANOVA. **(B)** Representation of the standard deviation of mRNA particle volume for each of the mRNA classes listed in **(A)** (n=3 images, all mRNA particles are represented from each image). **(C)** Covisualization of endogenous Me31B-GFP with *oskar* mRNA labeled with smFISH probes. **(D)** Overlap ratio calculations comparing the amount of Me31B-GFP condensates that overlap with *oskar* mRNA (n=24). **(E,F)** Same as **(C,D)** for *bicoid* mRNA (n=23). **(G,H)** Same as **(C,D)** for *nanos* mRNA (n=24). **(I,J)** Same as **(C,D)** for *me31B* mRNA (n=24). **(K,L)** Same as **(C,D)** for *cup* mRNA (n=24). **(M,N)** Same as **(C,D)** for *bruno1* mRNA (n=19). **(O,P)** Same as **(C,D)** for *armi* mRNA (n=24). **(Q,R)** Same as **(C,D)** for *glorund* mRNA (n=24). All images are XY projections of 5 optical Z slices of 0.3µm. Scale bars are 2µm. For all overlap ratio plots, each data point represents the average value of all Me31B-GFP condensates detected in an image. Significance was assessed using Mann-Whitney statistical tests. Error bars represent standard deviation. **** P < .0001.

Additionally, we calculated the SD of mRNA puncta volume for each species to compare the propensity of each mRNA to exist as multicopy mRNPs. mRNAs which exist primarily as single copies will have low volume SDs while mRNAs that exist in the cytoplasm both as single copies and as multi-copy puncta will have larger volume SDs (Fig. 3B). Unsurprisingly, all maternal mRNAs displayed volume SDs much larger than the non-P-body mRNAs. However, it was of note that the P-body component mRNAs seemed to behave independent of their class. *me31B* and *bruno1* mRNAs showed SDs much more comparable to the maternal mRNAs while *cup* mRNA showed a volume SD much closer to the non-P-body associated mRNAs. This may indicate that these mRNAs, while they are not putative P-body localized transcripts based on the literature, do display a propensity to form large multicopy mRNPs.

Based on these classifications, we first examined the localization of maternal mRNAs following 30 minutes of ER stress. Under basal conditions, P-bodies contain several *oskar* mRNA molecules; however, following DTT treatment, P-bodies appeared markedly enriched in *oskar* mRNA (Fig. 3C). Because the density of *oskar* mRNA puncti precluded reliable quantification of individual transcripts, we instead quantified the overlap volume ratio between Me31B-GFP labeled P-bodies and *oskar* mRNA. Using this approach, we observed an ∼4.99-fold increase in Me31B-GFP signal overlapping with *oskar* mRNA following ER stress (Fig. 3D). We next examined two additional maternal transcripts, *bicoid* and *nanos*, and observed similar trends. ER stress-induced an ∼3.33-fold increase in overlap between Me31B-GFP and *bicoid* mRNA and an ∼1.95-fold increase for *nanos* mRNA (Fig. 3E-H). Together, these results indicate that ER stress alters P-body composition by increasing the amount of maternal mRNAs enriched within these granules.

We next examined mRNAs encoding P-body components. Visualization of *me31B* mRNA following ER stress revealed a similar enrichment within P-bodies, with an ∼3.03-fold increase in overlap volume ratio between *me31B* mRNA and Me31B-GFP labeled P-bodies (Fig. 3I, J). This observation was notable because *me31B* mRNA is not typically considered a frequent P-body client based on prior studies and our own analysis (Fig. 3A). We then analyzed *cup* mRNA, which appeared largely excluded from P-bodies under basal conditions. Following DTT-induced ER stress, however, *cup* mRNA localized to P-bodies, exhibiting an ∼2.23-fold increase in overlap volume ratio (Fig. 3K, L). Finally, we examined *bruno1* mRNA and observed a pronounced enrichment within P-bodies, with an ∼11.49-fold increase in overlap ratio after ER stress (Fig. 3M, N). Bruno 1 protein is a well-characterized P-body-associated RNA-binding protein that binds target mRNAs and recruits Cup to transcripts, although its own transcript has not previously been reported as a strong P-body client (Nakamura et al. 2004). Notably, analysis of mRNA puncta volume SD revealed that *bruno1* mRNA exhibited a distribution similar to that of maternal transcripts known to localize to P-bodies (Fig. 3B, S3F). Together, these results suggest that ER stress promotes recruitment of mRNAs encoding P-body components to P-bodies, despite these transcripts not typically behaving as canonical P-body clients.

To determine whether this recruitment was specific, we next examined two transcripts not associated with P-bodies, *armi* and *glorund*. Both genes encode non-P-body associated mRNAs that are broadly expressed during oogenesis (Tomari et al. 2004; Kalifa et al. 2009). Consistent with previous reports and our Vantage plot analysis, these transcripts displayed large average distances from P-bodies and relatively small SD in puncta volume, indicating that they predominantly exist as single-copy puncta (Fig. 3A, B). In contrast to the transcripts described above, ER stress did not alter the association of *armi* or *glorund* mRNA with P-bodies (Fig. 3O, R). These observations indicate that ER stress does not drive indiscriminate transcript recruitment, but instead, promotes the selective enrichment of specific classes of mRNAs within P-bodies.

### mRNAs associated with P-bodies during ER stress are protected from degradation

ER stress activates the endoribonuclease Ire1, which initiates both XBP1 splicing and regulated Ire1-dependent decay (RIDD), a pathway that promotes widespread mRNA degradation to reduce ER load. In *D. melanogaster*, RIDD has been shown to be largely promiscuous, with transcript susceptibility to degradation governed primarily by proximity to the ER membrane rather than sequence features (Gaddam et al. 2013). Because nurse cells are highly enriched in ER membranes, this raised the possibility that many transcripts may be vulnerable to degradation during ER stress.

Given the selective recruitment of specific mRNAs to enlarged P-bodies following ER stress, we asked whether P-body association functions to protect transcripts from degradation. To broadly assess transcript fate, we quantified the abundance of the eight mRNAs by RT-qPCR following 30 minutes of ER stress. Notably, the two non-P-body-associated mRNAs, *armi* and *glorund*, showed a significant decrease in abundance (a ∼23.92% decrease for *armi* mRNA and a ∼22.84% decrease for *glorund* mRNA), indicating that ER stress is accompanied by mRNA degradation in nurse cells (Fig. 4A). In contrast, all mRNAs that were recruited to P-bodies during ER stress, including maternal mRNAs and P-body component mRNAs, showed increased overall abundances (*bcd* mRNA ∼14.46%, *nos* mRNA ∼31.18%, *osk* mRNA ∼16.63%, *cup* mRNA ∼21.72%, *bru1* mRNA ∼20.56%, and *me31B* mRNA ∼16.57%) (Fig. 4A).

**Figure 4:**
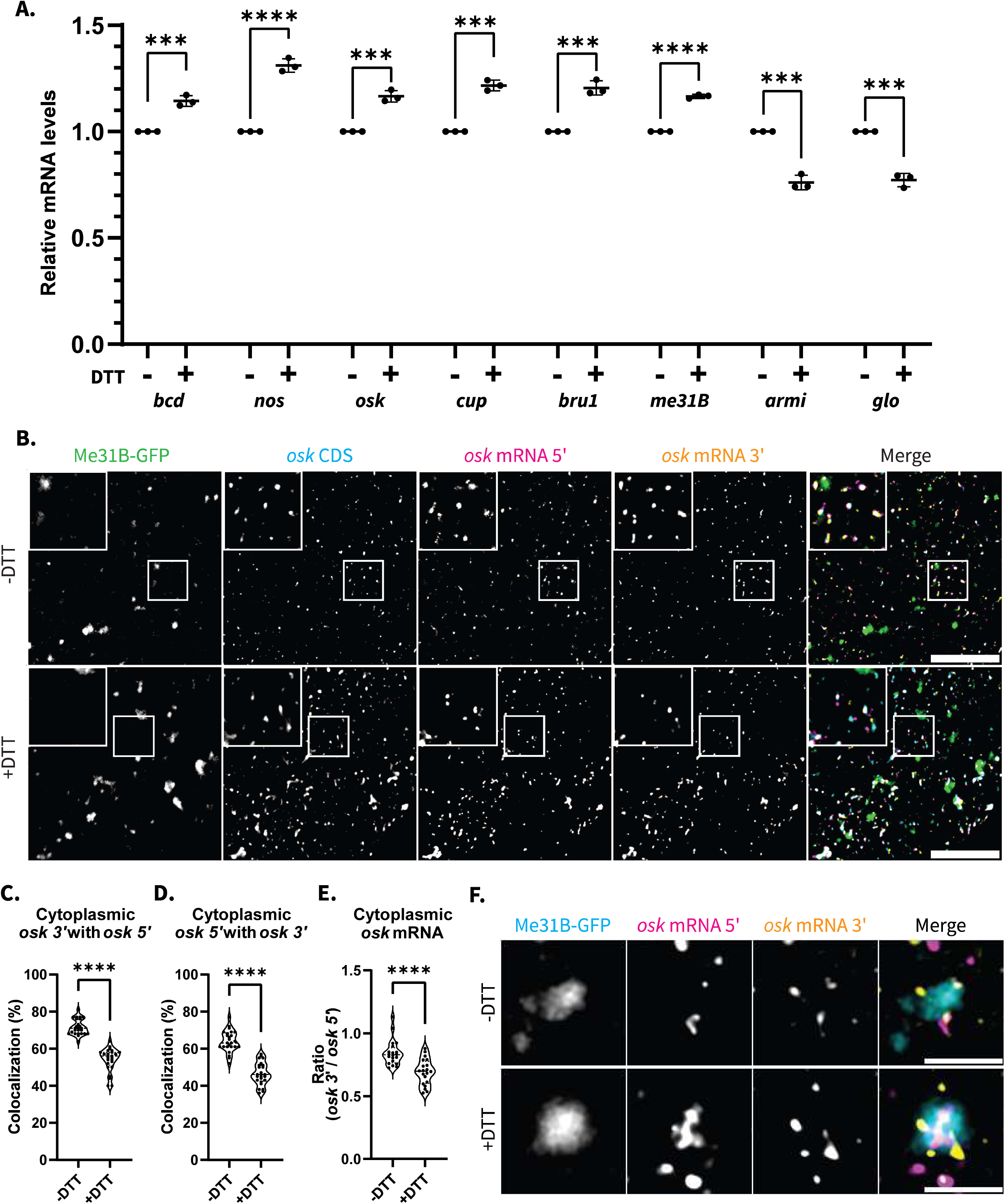
mRNAs associated with P-bodies during ER stress are protected from degradation. **(A)** RT-qPCR quantification of 8 mRNAs in −DTT and +DTT incubated egg chambers. Significance calculated with Welch’s t-test (n = 3). **(B)** Endogenously expressed Me31B-GFP covisualized with smFISH labeled oskar with differentially labeled CDS. 5′region and 3′ region. Images are XY projections of 5 optical Z slices of 0.3µm. Scale bars are 10µm. **(C)** Colocalization analysis between the *osk* 3′ and *osk* 5′ labeled puncta. Only puncta that also colocalized with the CDS were used in the calculations (n=25). **(D)** Colocalization analysis between the *osk* 5′ and *osk* 3′ labeled puncta. Only puncta that also colocalized with the CDS were used in the calculations (n=25). **(E)** Ratio of total *osk* 3′ puncta over total *osk* 5′ puncta (n=20). **(F)** Covisualization of endogenous Me31B-GFP with smFISH labeled *osk* 3′ and *osk* 5′ regions via STED. Images are XY projections of 5 optical Z slices of 0.22µm. Scale bar is 2µm. For all plots based on imaging, each data point represents the average value of all mRNAs detected in an image. Significance was assessed using Mann-Whitney statistical tests. Error bars represent standard deviation. **** P < .0001.

The surprising finding that transcript levels increased above control levels cannot be explained only by protection from degradation associated with ER stress as this does not occur under basal conditions. We hypothesized that the increase in the level of these mRNAs was due to enrichment of the transcripts within P-bodies where they would be protected from all background degradation even those they would be exposed to under basal conditions. Although these transcripts are not canonical targets of the mRNA decay machinery, it is possible that a fraction undergoes low-level “leaky” degradation under steady-state conditions. To address this at basal conditions, we evaluated the level of these mRNAs in the RNAi knockdown background of the key 5′ to 3′ exonuclease enzyme Pacman (Xrn1) to reduce the ‘leaky’ degradation. *pacman^RNAi^* was induced using the UAS/GAL4 system and the efficiency of the knockdown was confirmed via RT-qPCR (Fig. S4A). Consistent with our hypothesis, all the transcripts increased in abundance in this background, suggesting that under basal conditions there is low grade leaky degradation of some transcripts (Fig. S4B).

To determine if the increased levels of the P-body associated mRNAs indeed resulted from protection from degradation or possibly due to elevated transcription, we assessed transcriptional activity by measuring the volume of nascent transcription sites. Analysis of *oskar* transcription revealed no change in transcription site volume following ER stress (Fig. S4C, D). Similarly, transcription site volumes for *bicoid* and *nanos* were unchanged, indicating that increased maternal mRNA abundance does not arise from increased transcription, but instead results from increased protection from degradation (Fig. S4E-H).

We next asked whether during ER stress P-body associated maternal mRNAs are more protected from degradation versus cytoplasmic maternal mRNAs that are not within P-bodies. To assess cytoplasmic mRNA stability, we employed dual-color smFISH using probe sets differentially targeting the 5′ and 3′ ends of *oskar* mRNA and excluded from our analysis any signal that was colocalized with Me31B-GFP labeled P-bodies (Fig. 4B). To ensure that all 5′ and 3′ ends signals were representative of true *oskar* transcripts, we only quantified puncta which colocalized with the separately labeled *oskar* mRNA coding sequence. Protected or compact mRNAs exhibit reduced spatial separation between their 5′ and 3′ ends, resulting in increased probe colocalization (Adivarahan et al. 2018; Khong and Parker 2018). Following ER stress, cytoplasmic *oskar* mRNA showed a ∼24.43% decrease in 3′:5′ colocalization and a ∼27.52% decrease in 5′:3′ colocalization compared to control conditions, indicating decreased mRNA compaction and protection (Fig. 4C, D). Consistent with this interpretation, the ratio of 3′ to 5′ puncta decreased by ∼17.96% following ER stress, suggesting increased transcript degradation as there were more 5′ puncta than 3′ puncta in the cytoplasm (Fig. 4E). Together, this data indicates that maternal mRNAs that are not recruited to P-bodies during ER stress are more susceptible to cytoplasmic degradation. Following ER stress, a greater fraction of *oskar* transcripts localizes to P-bodies, accompanied by an overall increase in total *oskar* abundance. Despite this increase, we observe elevated degradation of *oskar* transcripts that remain outside of P-bodies. These findings support a model in which P-bodies function as protective compartments that selectively shield mRNAs from enhanced cytoplasmic decay during ER stress.

To further investigate the organization of *oskar* mRNA within P-bodies during ER stress, we utilized these same dual-color smFISH probes coupled with STED super resolution microscopy. Notably, under control conditions *oskar* 5′ and 3′ ends were colocalized within P-bodies but, following ER stress, the 5′ and 3′ ends segregated with the *oskar* 5′ ends localizing to the center of P-bodies and *oskar* 3′ ends localizing to the periphery of the condensates (Fig. 4F). Interestingly, this is consistent with previous work that showed that granule state can affect mRNA intracondensate organization (Ramat et al. 2024). Together, this data suggests that ER stress leads to a change in *oskar* mRNA stability with cytoplasmic particles being more prone to degradation and P-body associated particles reorganizing within the granules.

### Bruno 1 is upregulated during ER stress and is required for P-body remodeling

To determine whether increased abundance of P-body component mRNAs following ER stress was driven by transcriptional upregulation, we quantified transcription site volumes for *me31B*, *cup*, and *bruno1*. Transcription site analysis revealed no significant change in *me31B* or *cup* transcription following ER stress (Fig. S5A-D). Surprisingly, in contrast, *bruno1* transcription site volume increased by ∼80.12% after 30 minutes of ER stress, suggesting an overall increase in *bruno1* transcription (Fig. 5A, B). Furthermore, increased *bruno1* transcription resulted in elevated Bruno 1 protein levels detected via immunostaining following ER stress (Fig. 5C). Consistent with this observation, western blot analysis revealed an ∼47% increase in total Bruno 1 protein levels after 30 minutes of ER stress (Fig. 5D). Together, this data demonstrates that ER stress selectively induces *bruno1* transcription, resulting in increased Bruno 1 protein expression.

**Figure 5:**
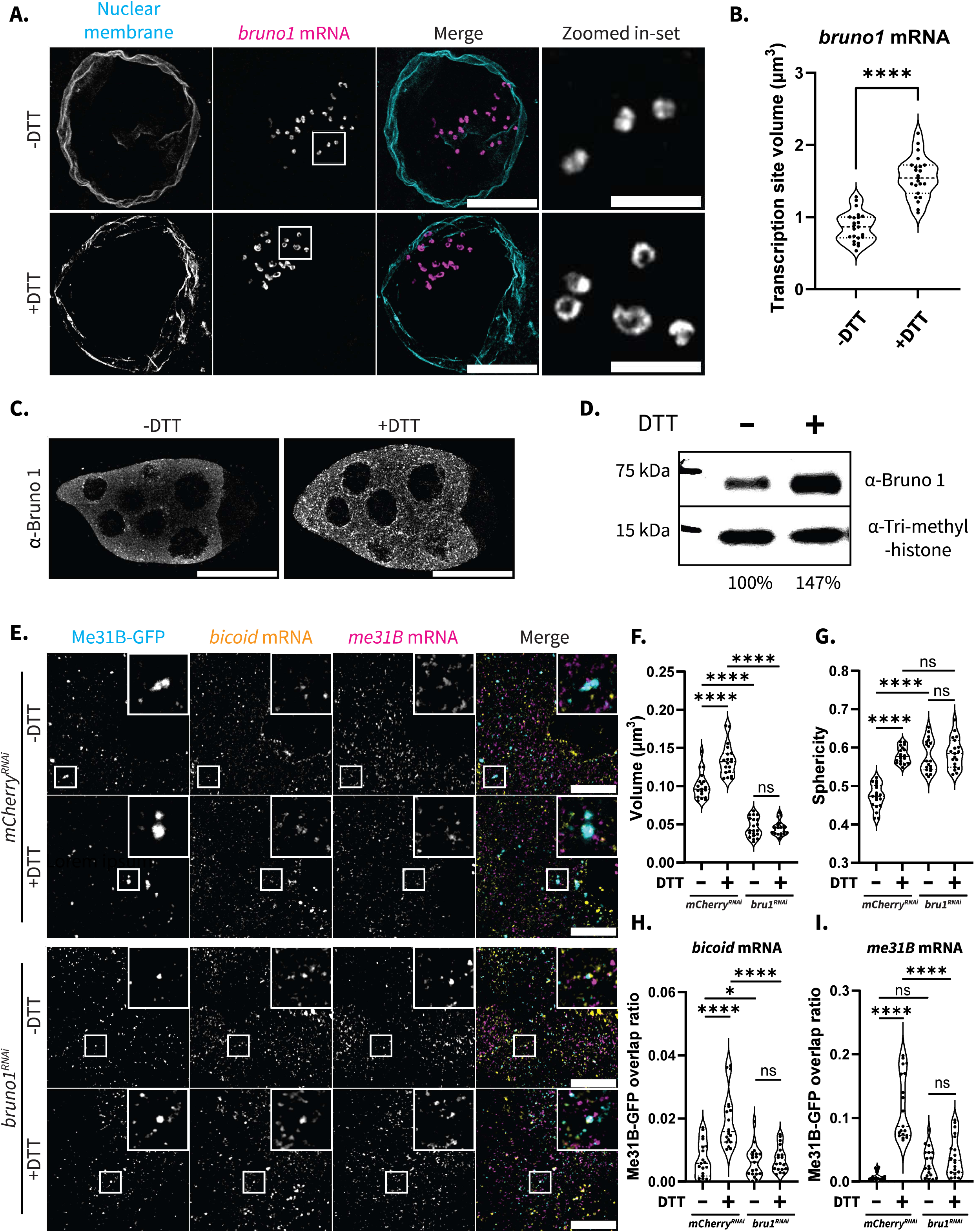
Bruno 1 is upregulated during ER stress and is required for P-body remodeling. **(A)** Covisualization of *bruno1* mRNA labeled with smFISH probes and the nuclear membrane visualized with wheat agglutinin staining. Images are XY projections of 15 optical Z slices of 0.3µm. Scale bars are 10µm. **(B)** Quantification of *bruno1* transcription site volume in −DTT and +DTT incubated egg chambers (n =23). **(C)** Visualization of immunolabeled Bruno 1 in −DTT and +DTT incubated egg chambers. Images are XY projections of 5 optical Z slices of 0.3µm. Scale bars are 100µm. **(D)** Western blot analysis of Bruno 1 in −DTT and +DTT incubated egg chambers. (Tri-methyl-Histone --loading control). **(E)** Covisualization of endogenously tagged Me31B-GFP with *bicoid* and *me31B* mRNA labeled with smFISH probes in −DTT and +DTT incubated *mCherry^RNAi^*(control) and *bruno1^RNAi^* egg chambers. Images are XY projections of 5 optical Z slices of 0.3µm. Scale bars are 20µm. **(F)** Volume quantifications for Me31B-GFP labeled condensates in **(E)** (n=19). **(G)** Sphericity quantifications for Me31B-GFP labeled condensates in **(E)** (n=19). **(H)** Overlap ratio calculations comparing the amount of Me31B-GFP condensates that overlap with *bicoid* mRNA in **(E)** (n=)19. **(I)** Overlap ratio calculations comparing the amount of Me31B-GFP condensates that overlap with *me31B* mRNA in **(E) (**n=19). For all plots, each data point represents the average value of all mRNAs and Me31B-GFP labeled condensates detected in an image. Significance was assessed using Mann-Whitney statistical tests. Error bars represent standard deviation. **** P < .0001.

Because Bruno 1 is a core P-body component that binds directly to mRNA and interacts with other P-body proteins, including Cup, we next asked whether Bruno 1 contributes to our previously observed ER stress-induced changes in P-body morphology (Bayer et al. 2023; Kim-Ha et al. 1995; Nakamura et al. 2004). To test this, we performed an RNAi-facilitated knockdown of Bruno 1 using the UAS/Gal4 system and visualized P-bodies using endogenously expressed Me31B-GFP (Fig. 5E). RNAi efficiency was confirmed via immunostaining (Fig. S5E). Because Bruno 1 knockdown egg chambers arrest at early developmental stages, we re-evaluated ER stress effects at stage 3 of oogenesis, as all our prior quantifications focused on mid-stage egg chambers.

Under basal conditions, Me31B-GFP marked P-bodies were ∼55.28% smaller in the *bruno1^RNAi^* background compared to control (*mCherry^RNAi^*) egg chambers, and ∼22.38% more spherical, indicating that Bruno 1 contributes to P-body morphology even in the absence of ER stress (Fig. 5F, G). Accompanying this reduction in condensate volume, we also observed an ∼2.77-fold increase in the overall number of P-bodies in the *bruno1^RNAi^* egg chambers compared to controls (Fig. S5F). Although we could not assess the overall level of Me31B protein via western blot analysis, as Bruno 1 knockdown ovaries do not develop past stage 4 of oogenesis and it is difficult to accurately compare protein levels at specific stages, immunofluorescence imaging suggests that overall Me31B levels remain the same (Fig. 5E). Together this data indicates that, in the absence of Bruno 1, P-bodies reorganize into an increased number of smaller, more spherical condensates.

Strikingly, whereas P-bodies in control egg chambers increased in volume by ∼22.87% and sphericity by ∼33.45% following ER stress, P-bodies in the Bruno 1 knockdown background showed no detectable change in morphology between basal and ER stress conditions (Fig. 5E-G). This indicates that Bruno 1 is necessary for ER stress-induced P-body enlargement and confirms that P-bodies respond to ER stress at multiple stages of development.

We next asked whether Bruno 1 is also required for functional changes in P-body composition during ER stress. To assess this, we quantified the overlap volume ratio between Me31B-GFP labeled P-bodies and representative P-body associated mRNAs (*bicoid* and *me31B*) to assess mRNA association with P-bodies. We chose not to use *oskar* as our maternal mRNA representative as Bruno 1 has been shown to directly dictate its post-transcriptional regulation and this may make it difficult to parse changes at the condensate level (Kim-Ha et al. 1995; Snee et al. 2008). In control egg chambers, ER stress led to an ∼14.42-fold increase in Me31B-GFP overlap ratio with *bicoid* mRNA and an ∼2.77-fold increase in overlap ratio with *me31B* mRNA (Fig. 5E). In contrast, in the *bruno1^RNAi^* background, ER stress failed to significantly increase the enrichment of either mRNA in P-bodies (Fig. 5H, I). Thus, Bruno 1 is required, not only for ER stress-induced changes in P-body morphology, but also for the consequent recruitment of mRNAs to P-bodies during the ER stress response.

### ATF4 acts upstream of Bruno 1 during ER stress response

Since we found that increased Bruno 1 levels are required for ER stress-induced P-body remodeling, we sought to decipher the underlying mechanism of how ER stress signaling leads to *bruno1* upregulation. The UPR activates three major transcriptional arms leading to the nuclear localization of three transcription factors, XBP1, ATF4, and ATF6 (Walter and Ron 2011). These three proteins are highly conserved, and in *D. melanogaster*, the ATF4 gene name is *cryptocephal* (*crc*) (Hewes et al. 2000). To determine which pathway is required for P-body remodeling, we examined Me31B-GFP labeled P-bodies in *XBP1^RNAi^*and *ATF4*/*crc^RNAi^* backgrounds (Fig. 6A). Knockdown efficiency was confirmed via western blotting for *XPB1^RNAi^* and RT-qPCR for *ATF4/crc^RNAi^* (Fig. S6A, B).

**Figure 6:**
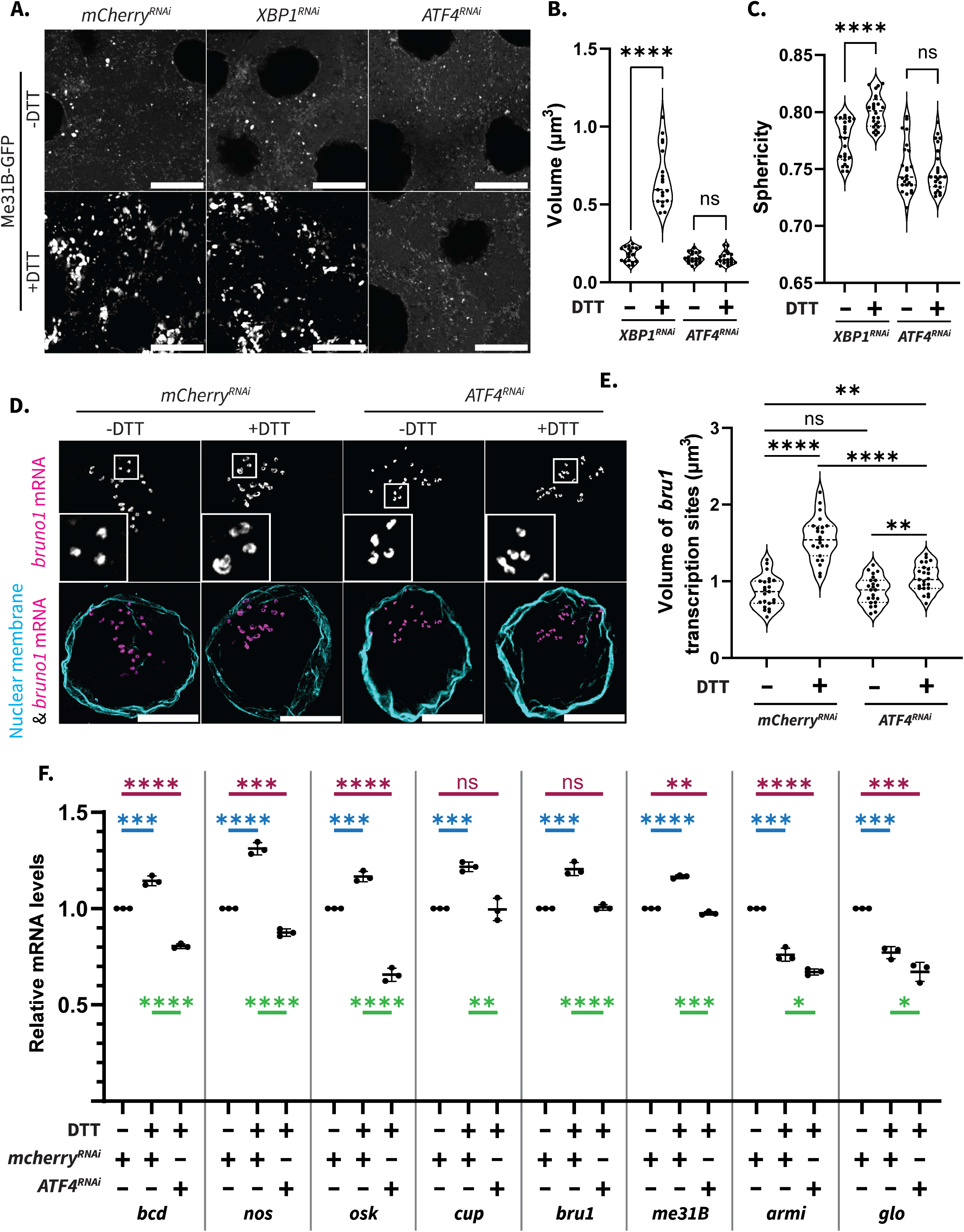
ATF4 acts upstream of Bruno 1 during ER stress response. **(A)** Endogenously labeled Me31B-GFP visualized in *mCherry^RNAi^, XBP1^RNAi^,* and *ATF4/crc^RNAi^* egg chambers incubated in −DTT and +DTT. Images are XY projections of 5 optical Z slices of 0.3µm. Scale bars are 20µm. **(B)** Volume quantifications for Me31B-GFP condensates in **(A)** (n=25). **(C)** Sphericity quantifications for Me31B-GFP condensates in **(A)** (n=24). **(D)** *bruno1* mRNA labeled with smFISH probes visualized with wheat agglutinin nuclear membrane stain in *mCherry^RNAi^* and *ATF4/crc^RNAi^* egg chambers incubated in −DTT and +DTT. Images are XY projections of 15 optical Z slices of 0.3µm. Scale bars are 10µm. **(E)** Quantification of *bruno1* transcription site volume in **(D)** (n =20). **(F)** RT-qPCR quantification of 8 mRNAs in *mCherry^RNAi^* and *ATF4/crc^RNAi^* egg chambers incubated in −DTT and +DTT. Significance calculated with Welch’s t-test (n = 3). For all plots based on imaging, each data point represents the average value of all Me31B-GFP labeled condensates or mRNAs detected in an image. Significance was assessed using Mann-Whitney statistical tests. Error bars represent standard deviation. **** P < .0001.

In control egg chambers, ER stress-induced robust enlargement and increased sphericity of Me31B-GFP marked P-bodies after 30 minutes (Fig. 6B, C). A similar response was observed in the *XBP1^RNAi^* background with an ∼4.25-fold increase in P-body volume and an ∼3.35% increase in P-body sphericity, indicating that XBP1 is dispensable for ER stress-induced P-body remodeling (Fig. 6B, C). In contrast, P-bodies in the *ATF4/crc^RNAi^* background failed to undergo any detectable morphological changes following ER stress, phenocopying the *bruno1^RNAi^* egg chambers (Fig. 6B, C). These results suggest that P-body remodeling occurs downstream of the ATF4 arm of the ER stress response.

To test whether ATF4 is required for ER stress-induced increased *bruno1* transcription, we quantified *bruno1* transcription sites in the *ATF4*/*crc^RNAi^*background. Under basal conditions, *bruno1* transcription was comparable between control and ATF4 knockdown egg chambers. However, following ER stress, *bruno1* transcription increased only modestly, ∼12.63% in the absence of ATF4, compared to ∼81.13% in control egg chambers (Fig. 6D, E). Consistent with this result, immunostaining revealed no appreciable increase in Bruno 1 protein levels following ER stress in the *ATF4/crc^RNAi^* background (Fig. S6C). Together, this data indicates that ER stress-induced upregulation of Bruno 1 is largely dependent on ATF4 activity.

Finally, we asked whether loss of ATF4 affected the fate of the eight representative mRNAs during ER stress. To assess this, we quantified transcript abundance following ER stress in the *ATF4*/*crc^RNAi^*background. As in controls, non-P-body-associated mRNA levels decreased following ER stress (*armi* decreased by ∼32.88% and *glorund* decreased by ∼32.83%) (Fig. 6F). In contrast, maternal mRNAs that were protected and increased in abundance in control egg chambers, instead decreased significantly in the absence of ATF4 (*bicoid* mRNA by ∼19.49%, *nanos* mRNA by ∼12.47%, and *oskar* mRNA by ∼34.36%). Similarly, P-body component mRNAs failed to increase in abundance following ER stress in the *ATF4*/*crc^RNAi^*background (Fig. 6F). Thus, loss of ATF4 abolishes both P-body remodeling and selective mRNA protection during ER stress.

### Bruno 1 overexpression is sufficient to drive P-body remodeling and rescues morphology in the ATF4 knockdown background

Because loss of ATF4 abolished both ER stress-induced P-body remodeling and Bruno 1 upregulation, we next asked whether increasing Bruno 1 levels alone is sufficient to restore condensate morphological response following ER stress. To test this, we again used the UAS/GAL4 system to specifically upregulate transgenic Bruno 1-GFP expression in the female germline. Previous studies have demonstrated that this overexpression line is viable (Nikonova et al. 2024). Immunofluorescence analysis confirmed that this approach resulted in elevated Bruno 1 protein levels (Fig. S7A, A’). Under basal conditions, overexpression of Bruno 1-GFP in *mCherry^RNAi^* control egg chambers led to a marked change in P-body morphology, with Me31B-GFP labeled P-bodies exhibiting an ∼58.32% increase in volume and an ∼5.12% increase in sphericity (Fig. 7A-C). Notably, a similar increase in P-body volume and sphericity was observed in the *ATF4*/*crc^RNAi^* egg chambers, indicating that elevated Bruno 1 levels are sufficient to promote P-body remodeling independent of ATF4 activity.

**Figure 7:**
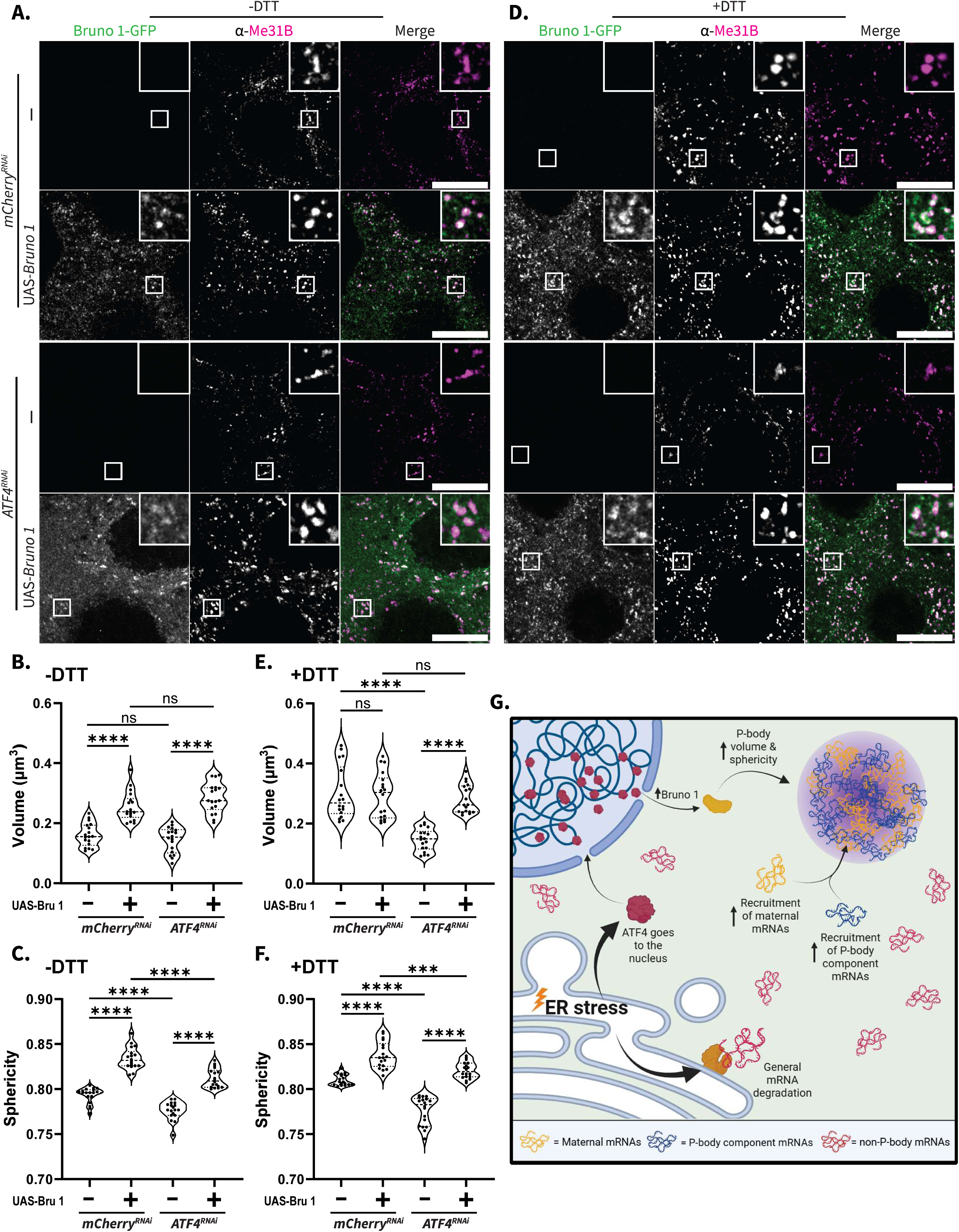
Bruno 1 overexpression is sufficient to drive P-body remodeling and rescues morphology in the ATF4 knockdown background. **(A)** Covisualization of transgenic Bruno 1-GFP and immunolabeled Me31B in *mCherry^RNAi^* and *ATF4/crc^RNAi^* egg chambers incubated in Schneider’s media (-DTT). Images are XY projections of 5 optical Z slices of 0.3µm. Scale bars are 20µm. **(B)** Volume quantifications for Me31B condensates in **(A)** (n=18). **(C)** Sphericity quantifications for Me31B condensates in **(A)** (n=18). **(D)** Covisualization of transgenic Bruno 1-GFP and immunolabeled Me31B in *mCherry^RNAi^* and *ATF4/crc^RNAi^* egg chambers incubated in Schneider’s media supplemented with DTT (+DTT). Images are XY projections of 5 optical Z slices of 0.3µm. Scale bars are 20µm. **(E)** Volume quantifications for Me31B condensates in **(D)** (n=18). **(F)** Sphericity quantifications for Me31B condensates in **(D)** (n=18). **(G)** Proposed mechanism of how ER stress triggers P-body response. For all plots, each data point represents the average value of all Me31B-labeled condensates or mRNAs detected in an image. Significance was assessed using Mann-Whitney statistical tests. Error bars represent standard deviation. **** P < .0001.

We next examined whether further increases in Bruno 1 levels could augment P-body remodeling during ER stress (Fig. 7D). In the *mCherry^RNAi^* background, Bruno 1 overexpression increased P-body sphericity but did not further increase P-body volume following ER stress, suggesting that P-body volume may be constrained by a threshold level of Bruno 1 beyond which there is no additional change in size (Fig. 7E, F). Alternatively, there may be a maximum sustainable P-body volume in this tissue.

In contrast, in the *ATF4*/*crc^RNAi^* egg chambers, Bruno 1 overexpression restored P-body volume to levels comparable to control egg chambers following ER stress and led to an even greater increase in P-body sphericity (Fig. 7E, F). Together, these results demonstrate that Bruno 1 upregulation is sufficient to drive P-body remodeling and can compensate for the loss of ATF4 during the ER stress response.

## Discussion

Cytoplasmic condensates such as P-bodies have long been associated with mRNA repression and decay, yet how their properties are actively regulated during cellular stress remains unclear (Luo et al. 2018; Standart and Weil 2018; Eulalio et al. 2007). Early work in yeast and mammalian systems established P-bodies as dynamic assemblies whose size and number respond to translational status and stress conditions (Brengues 2005; Teixeira et al. 2005; Parker and Sheth 2007). More recent studies have emphasized that P-bodies are compositionally heterogeneous, and exhibit liquid-like behaviors shaped by multivalent RNA:protein interactions (Banani et al. 2016; Sankaranarayanan et al. 2021). Our findings extend this framework by demonstrating that ER stress rapidly remodels P-body morphology and internal organization in *D. melanogaster* nurse cells, revealing P-bodies as early and responsive components of the ER stress response rather than passive byproducts of translational repression.

Several studies have suggested that stress granules and P-bodies can exchange components and often respond sequentially to stress, with stress granules typically forming after prolonged translational arrest (Kedersha et al. 2005; Riggs et al. 2020). Consistent with this view, we observe pronounced P-body remodeling within 30 minutes of ER stress induction, in the absence of detectable stress granule formation. This temporal separation suggests that P-bodies may act as an early buffering compartment during ER stress, potentially stabilizing select transcripts before engagement of later stress granule mediated responses. Such a model aligns with recent proposals that distinct condensates fulfill temporally and functionally specialized roles during stress adaptation (Kroschwald et al. 2018; Choi et al. 2025).

Mechanistically, our data identifies transcriptional upregulation of the RNA-binding protein Bruno 1 as a key driver of ER stress-induced P-body remodeling (Fig. 7G). Previous work has established Bruno1 as a translational repressor essential for maternal mRNA regulation during oogenesis and as a component of P-bodies (Webster et al. 1997; Chekulaeva et al. 2006; Bayer et al. 2023). By showing that ER stress induces upregulation of *bruno1* transcription in an ATF4-dependent manner, and that loss of Bruno 1 abolishes P-body compositional remodeling, we uncover a direct link between stress-responsive transcriptional programs and condensate regulation. This supports emerging models in which changes in the abundance of specific scaffold or client proteins can reshape condensate properties, rather than condensate behavior being governed solely by global changes in translation or RNA availability (Sankaranarayanan et al. 2021; Banani et al. 2016; Bayer et al. 2023).

Functionally, we find that P-body remodeling during ER stress is associated with selective recruitment and stabilization of specific mRNAs, including maternal transcripts and P-body component mRNAs, while non-associated transcripts are degraded. This is consistent with transcriptomic data that shows that upon arsenite stress, the composition of P-bodies drastically changes (Matheny et al. 2019). ER stress activates Ire1-mediated RNA decay through the RIDD pathway, which in *D. melanogaster* has been shown to act broadly on ER-proximal transcripts with limited sequence specificity (Gaddam et al. 2013). In this context, selective sequestration of mRNAs into P-bodies may provide a spatial mechanism to shield essential transcripts from general degradation. Our observation that P-body associated mRNAs exhibit increased abundance without increased transcription supports a protective, rather than degradative, role for P-bodies under these conditions, consistent with prior observations that P-bodies can function as sites of mRNA storage (Standart and Weil 2018; M.-D. Lin et al. 2008).

Finally, our data places P-body remodeling downstream of the ATF4, but independent of XBP1, highlighting functional specialization within ER stress signaling pathways. While XBP1 primarily regulates transcriptional programs that expand ER folding capacity, ATF4 has been implicated in broader adaptive responses, including glycolysis, amino acid metabolism and stress resistance (Ji Eun Lee et al. 2015; Read and Schröder 2021; Walter and Ron 2011). Notably, previous work has found that ATF4 induces increased transcription of 4E-BP, a protein involved in post-transcriptional translation repression (Kang et al. 2017). Our findings extend the scope of ATF4 signaling to include regulation of RNA:protein condensates and post-transcriptional mRNA fate. Because ATF4 signaling can also be activated by the integrated stress response (ISR), these observations raise the possibility that P-body remodeling may participate in stress adaptation beyond ER stress (Pakos-Zebrucka et al. 2016). More broadly, this work suggests that transcriptional stress responses can directly reconfigure the physical organization of the cytoplasm to selectively preserve key gene products during stress, providing a new conceptual link between stress signaling, condensate biology, and mRNA regulation.

## Methods and Protocols

### Fly husbandry

*Drosophila melanogaster* stocks were maintained on standard cornmeal agar food at 25°C. Female flies were put in grape vials and fed yeast paste 2-3 days prior to dissection. Fly stocks obtained from Bloomington *Drosophila* Stock Center: UAS-*mCherry^RNAi^* (BL #35785), UAS-*bruno1^RNAi^* (BL #35394), *Me31B-GFP* (BL #51530), *tub-a*(V37)-Gal4 (BL #7063), *osk-a*(VP16)-Gal4 (BL #44241), UAS-*XBP1^RNAi^ ^(^*BL #36755), UAS- *crc^RNAi^* (ATF4) (BL #80388). Kyoto *Drosophila* Stock Center: *Cup-YFP* (DGRC 115-161). Vienna *Drosophila* Resource Center: *Rin-GFP* (VDCR #318907). *Tral-RFP* was a kind gift from Dr. D. St. Johnston (Gurdon Institute at the University of Cambridge), and UAS-*Bruno 1-GFP* was a kind gift from Dr. M. L. Spletter (University of Missouri-Kansas City). UAS-*mCherry^RNAi^* (BL #35785) was used as a control in all RNAi experiments to account for effects of activated RNAi machinery. All RNAi lines were driven by *tub-a*(V37)-Gal4 (BL #7063) except for UAS-*Bruno-GFP* experiments which required a Gal4 driver on a different chromosome, so *osk-a*(VP16)-Gal4 (BL #44241) was used.

### Immunofluorescence staining of *D. melanogaster* egg chambers

Ovaries were dissected and fixed in 2% PFA in PBS for 10 minutes at room temperature. Fixed egg chambers were washed three times for 10 minutes each in PBST (PBS containing 0.3% Triton X-100), then permeabilized and blocked for 2 hours in PBS containing 1% Triton X-100 and 1% BSA. Samples were incubated with primary antibodies overnight at room temperature with gentle rocking, followed by three 10-minute washes in PBST. Secondary antibody incubation was carried out using fluorescently labeled antibodies (1:1000; DyLight 550 & 650; ThermoFisher) for 2 hours at room temperature, followed by three additional 10-minute washes in PBST. Samples were mounted in RapiClear (SUNjin Lab) mixed with Aqua-Poly/Mount (Polysciences) at a 75:25 ratio for imaging. The following antibodies and stains were used: mouse anti-Me31B (1:3000), a generous gift from Dr. A. Nakamura (Institute of Molecular Embryology and Genetics, Kumamoto University), Rabbit anti-eIF4G (1:200) a kind gift from Dr. E. Izaurralde (Max Planck Institute for Developmental Biology), Rabbit anti-Bruno 1 (1:10,000) kind gift from Dr. P. M. Macdonald (University of Texas Austin).

### smFISH labeling

Single-molecule fluorescence *in situ* hybridization (smFISH) was performed following the protocol described by (Bayer et al. 2015), with minor modifications. Ovaries were dissected and fixed in 4% PFA in PBS for 10 minutes at room temperature. Fixed egg chambers were washed three times for 10 minutes each in 2x SSC, then pre-hybridized with a 15-minute wash in 2x SSC containing 10% formamide. Samples were then incubated overnight at 37 °C with smFISH probes (1:50) and a nuclear membrane stain, wheat germ agglutinin conjugated to CF405S (1:50; Biotium). Following hybridization, egg chambers were washed three times for 10 minutes each in pre-warmed 2x SSC with 10% formamide at 37 °C and mounted in ProLong Diamond Antifade Mountant (Life Technologies) for imaging. The following probe sets were used: *nanos* mRNA labeled with 48 Quasar 670 probes, *bicoid* mRNA labeled with 48 Quasar 570 probes, *me31B-GFP* mRNA labeled with 30 EGFP recognizing Quasar 670 probes, *cup* mRNA labeled with 48 Cal Fluor Red 590 probes, *armi* mRNA labeled with 48 Cal Fluor Red 590 probes, *glorund* mRNA labeled with 35 Quasar 590 probes, *oskar* mRNA CDS labeled with 48 Quasar 670 probes, *oskar* 5′ mRNA labeled with 46 Quasar 570 probes, *oskar* 3′ mRNA labeled with 45 Cal Fluor Red 610 probes, *bruno1* mRNA labeled with 48 Quasar 670 probes. Probes were made by Biosearch technologies.

### ER stress induction

Egg chambers were dissected directly into Schneider’s media alone (-DTT) or supplemented with 5 mM DTT (+DTT), or 1 µM Thapsigargin (+Thapsigargin). Following 30 minutes incubation, egg chambers were rinsed with 1x PBS and then fixed and either mounted as previously described or used for smFISH or immunofluorescence labeling as previously described.

### Microscopy

All imaging was performed using a Leica TCS SP8 Laser Scanning Confocal Microscope equipped with a white light laser (470-670 nm), a 405 nm solid-state laser, and a continuous-wave STED 660 nm high-intensity laser. For confocal imaging, a 63x/1.4 NA oil immersion objective and a and a 100x/1.4 NA oil immersion objective were used. Optical Z-sections were acquired at 0.3 µm intervals. For STED imaging, a 100x/1.4 NA oil immersion objective was used with a zoom factor of 5x, and optical Z-sections were acquired at 0.22 µm intervals. STED samples were prepared from 25 µm ovary cryosections. All images were acquired using an automated XYZ piezoelectric stage and saved as 16-bit files. Image acquisition was carried out using Leica LAS X software.

### Tissue preparation for sectioning

Ovaries were dissected directly into 4% PFA in 1x PBS and fixed for 10 minutes at room temperature. Samples were then washed three times for 10 minutes each in PBST (PBS with 0.1% Triton X-100), followed by a 5-minute wash in 0.1 M glycine, pH 3.0. After fixation and washing, ovaries were incubated overnight at 4 °C in 30% sucrose. The following day, samples were embedded in O.C.T. Compound (Tissue-Tek) and flash frozen prior to sectioning. Cryosections of 25 µm thickness were prepared using a cryotome and stored at −80 °C until use.

### Imaging analysis

Identical acquisition settings were used for all control and experimental samples to ensure comparability. For each condition, images were obtained from similar stage egg chambers from three independent experiments, each prepared from a separate fly cross. All raw images were deconvolved prior to analysis using Leica’s Lightning deconvolution module. Image processing was carried out using consistent batch parameters across all samples. Quantitative analyses were performed using Imaris Microscopy Image Analysis software (Oxford Instruments). Condensates, proteins, and mRNAs were detected using the Imaris “Surface” module, which enables object-based segmentation and object-to-object spatial analysis. Colocalization was determined using the “Surface-Surface” analysis function, where objects with a shortest edge-to-edge distance of less than 0 µm were classified as colocalized. All statistical analyses were conducted using Mann-Whitney statistical tests in GraphPad Prism 8 (GraphPad Software). Figures were assembled and image adjustments applied uniformly using Fiji/ImageJ (NIH) (Schindelin et al. 2012).

### Western blot analysis

For each genotype, ten ovaries were dissected directly into 95 µL of 2x Laemmli Sample Buffer (Bio-Rad) supplemented with 5 µL β-mercaptoethanol (BME) and immediately subjected to mechanical lysis. Samples were heated at 95 °C for 10 minutes and then centrifuged at 10,000x g for 10 minutes at 4 °C. Clarified lysates were loaded onto 10% SDS-PAGE acrylamide gels for electrophoretic separation. Primary antibodies used included mouse anti-Me31B (1:3000), generously provided by Dr. A. Nakamura (Institute of Molecular Embryology and Genetics, Kumamoto University); rabbit anti-Tri-methyl-Histone H3 (C42D8) (1:150,000; Cell Signaling Technology); rabbit anti-Bruno (1:10,000) a generous gift from Dr. P. M. MacDonald (University of Texas at Austin); rabbit anti-XBP1 (PB9463) (1:500; Boster Bio). Bands were detected using TrueBlot ULTRA secondary antibodies (anti-mouse and anti-rabbit IgG HRP, 1:50,000; Rockland) and visualized with SuperSignal West Femto Maximum Sensitivity Substrate (ThermoFisher Scientific).

### RNA isolation and RT-qPCR

Whole ovaries were dissected into ice-cold PBS (4 °C) and immediately subjected to mechanical lysis in TRIzol reagent (ThermoFisher Scientific) for total RNA extraction. RNA was precipitated and washed with ethanol and resuspended in RNase-free water. RNA concentration and purity were assessed prior to downstream applications. Reverse transcription was performed using 2.5 µg of total RNA with the Superscript IV First-Strand Synthesis Kit (Life Technologies) according to the manufacturer’s instructions. Primers were designed using the DRSC FlyPrimerBank and synthesized by Integrated DNA Technologies. Quantitative PCR reactions were carried out on a Roche LightCycler 480 system (Roche Molecular Systems, Inc.). Each 10 µL reaction contained 1 µL of cDNA template, 4 µL of 10 µM primer mix, and 5 µL of SYBR Green I Master Mix (Roche Diagnostics). Reactions were performed in triplicates. Relative expression levels were calculated and normalized using Rp-49, and statistical significance was evaluated using a two-tailed unpaired Welch’s t-test.

### Graphics

BioRender was used to prepare Figures 1A and 7G.

## Acknowledgments

We thank Dr. A. Nakamura, (Riken Center for Developmental Biology) and Dr. P. M. MacDonald (University of Texas at Austin) for the kind gifts of antibodies. We extend thanks to Dr. D. St Johnston (University of Cambridge) and Dr. M. L. Spletter (University of Missouri-Kansas City) for the kind gifts of *D. melanogaster* lines. We thank the BDSC Indiana and DGRC Kyoto for providing *D. melanogaster* lines, as well as the TRiP at Harvard Medical School (NIH/NIGMS RO1-GM084947) for the transgenic RNAi stocks. We thank the Bioimaging Facility at Hunter College for access to the Leica TCS SP8 and Imaris -- Image Analysis Software. We thank Dr. P. Feinstein (Hunter College) for allowing us to use the Roche Light-cycler instrument.

## Funding

This work was supported by the National Institute of Health (1SC1GM135132) and the National Science Foundation instrumentation award (1919829) to D. P. B.

## Data Availability

This study includes no data deposited in external repositories.

## Notes

### Competing Interest Statement

The authors have declared no competing interest.

