## Supplemental Figures for "ATF4-dependent upregulation of Bruno 1 remodels P-bodies to selectively protect mRNAs during ER stress throughout *Drosophila melanogaster* oogenesis"

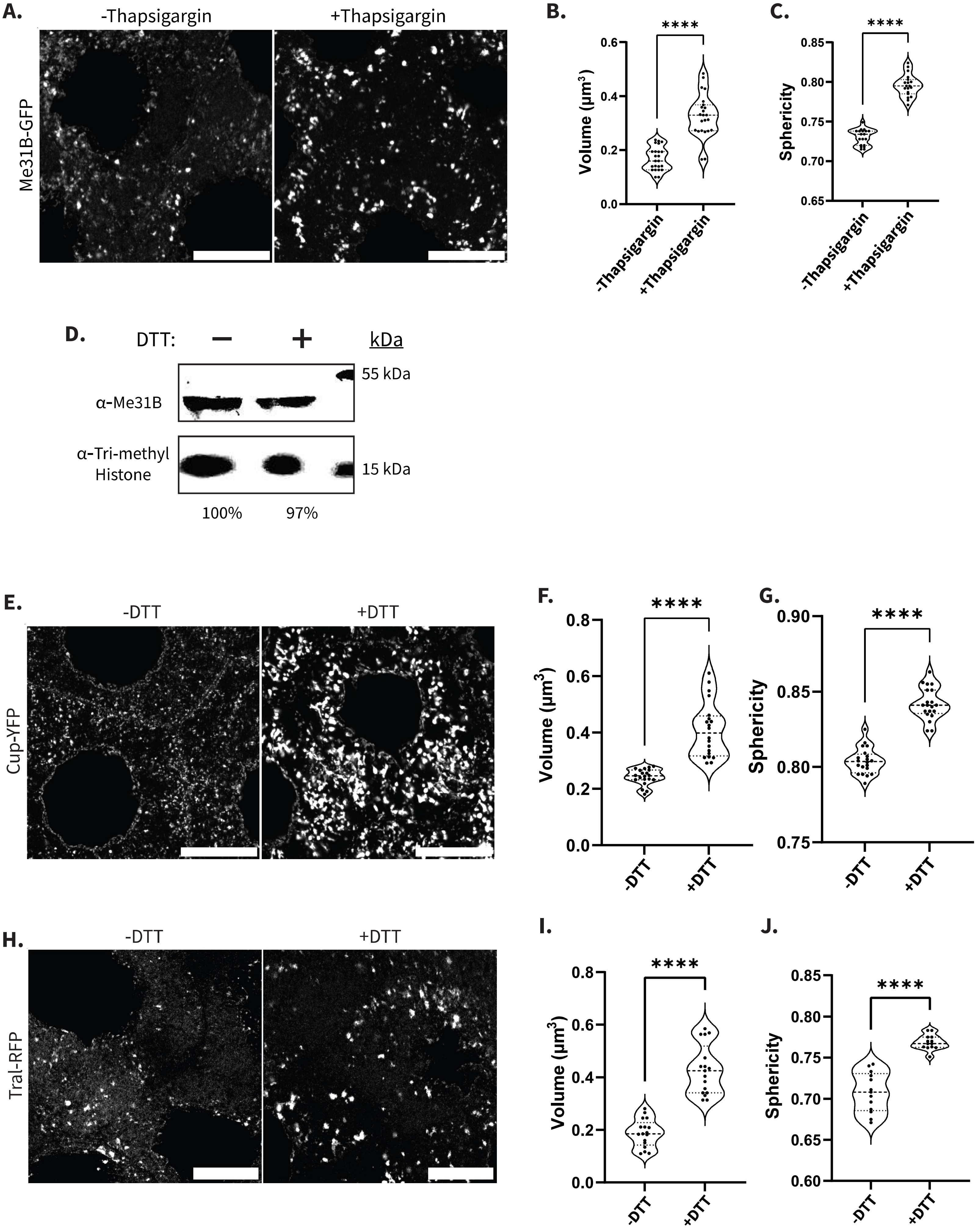
A.
-Thapsigargin
+Thapsigargin
Me31B-GFP
B.
****
Volume (µm³)
0.0
0.2
0.4
0.6
-Thapsigargin
+Thapsigargin
C.
****
Sphericity
0.65
0.70
0.75
0.80
0.85
-Thapsigargin
+Thapsigargin
D.
DTT:
−
+
kDa
55 kDa
α-Me31B
α-Tri-methyl
Histone
15 kDa
100%
97%
E.
-DTT
+DTT
Cup-YFP
F.
****
Volume (µm³)
0.0
0.2
0.4
0.6
0.8
-DTT
+DTT
G.
****
Sphericity
0.75
0.80
0.85
0.90
-DTT
+DTT
H.
-DTT
+DTT
Tral-RFP
I.
****
Volume (µm³)
0.0
0.2
0.4
0.6
0.8
-DTT
+DTT
J.
****
Sphericity
0.60
0.65
0.70
0.75
0.80
0.85
-DTT
+DTT

Figure S1

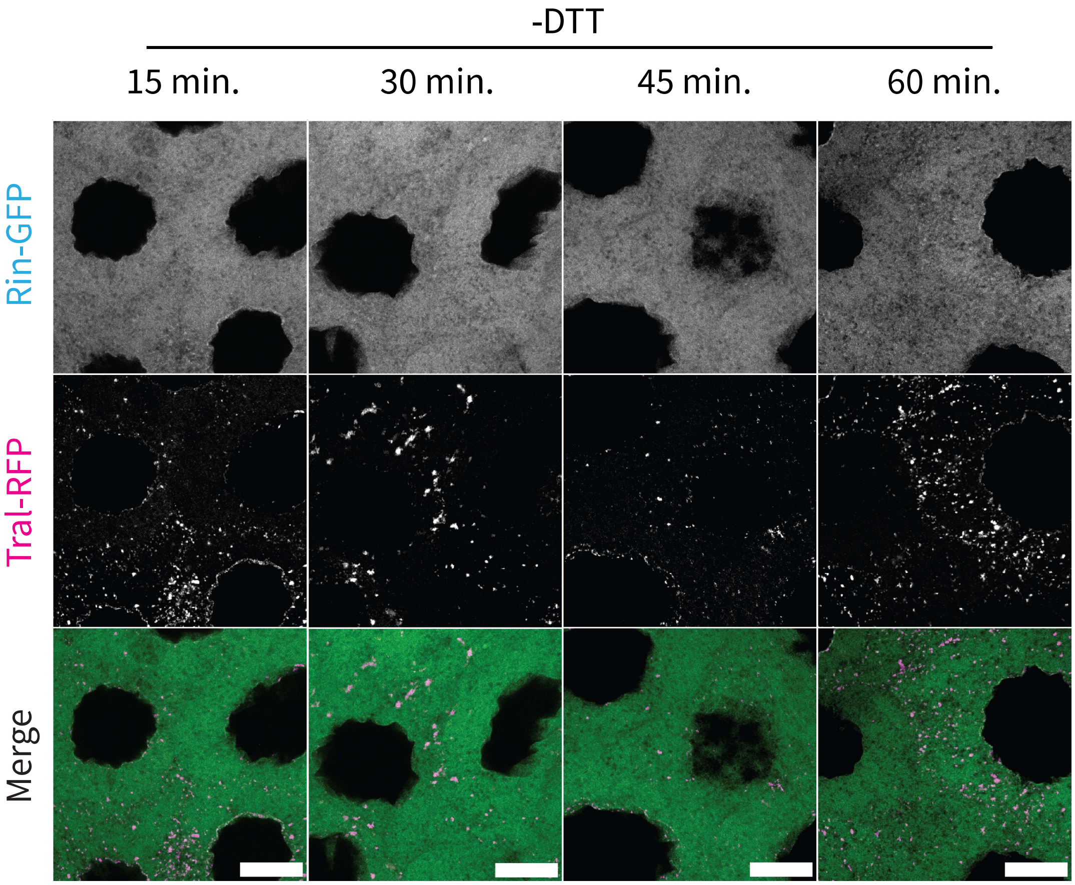


Figure S2

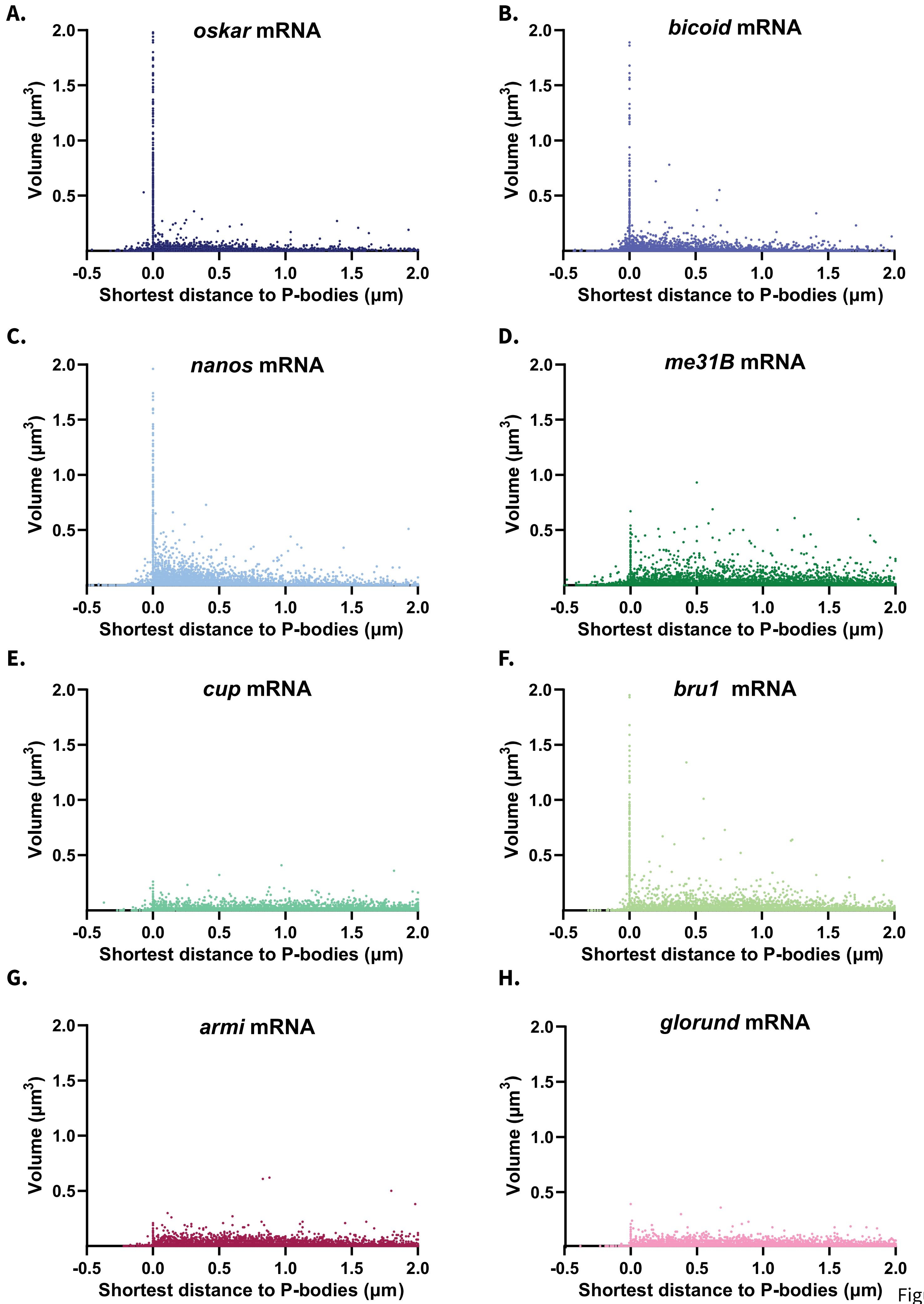

A.
oskar mRNA
B.
bicoid mRNA
C.
nanos mRNA
D.
me31B mRNA
E.
cup mRNA
F.
bru1 mRNA
G.
armi mRNA
H.
glorund mRNA
Volume (μm3)
Shortest distance to P-bodies (μm)


Figure S3

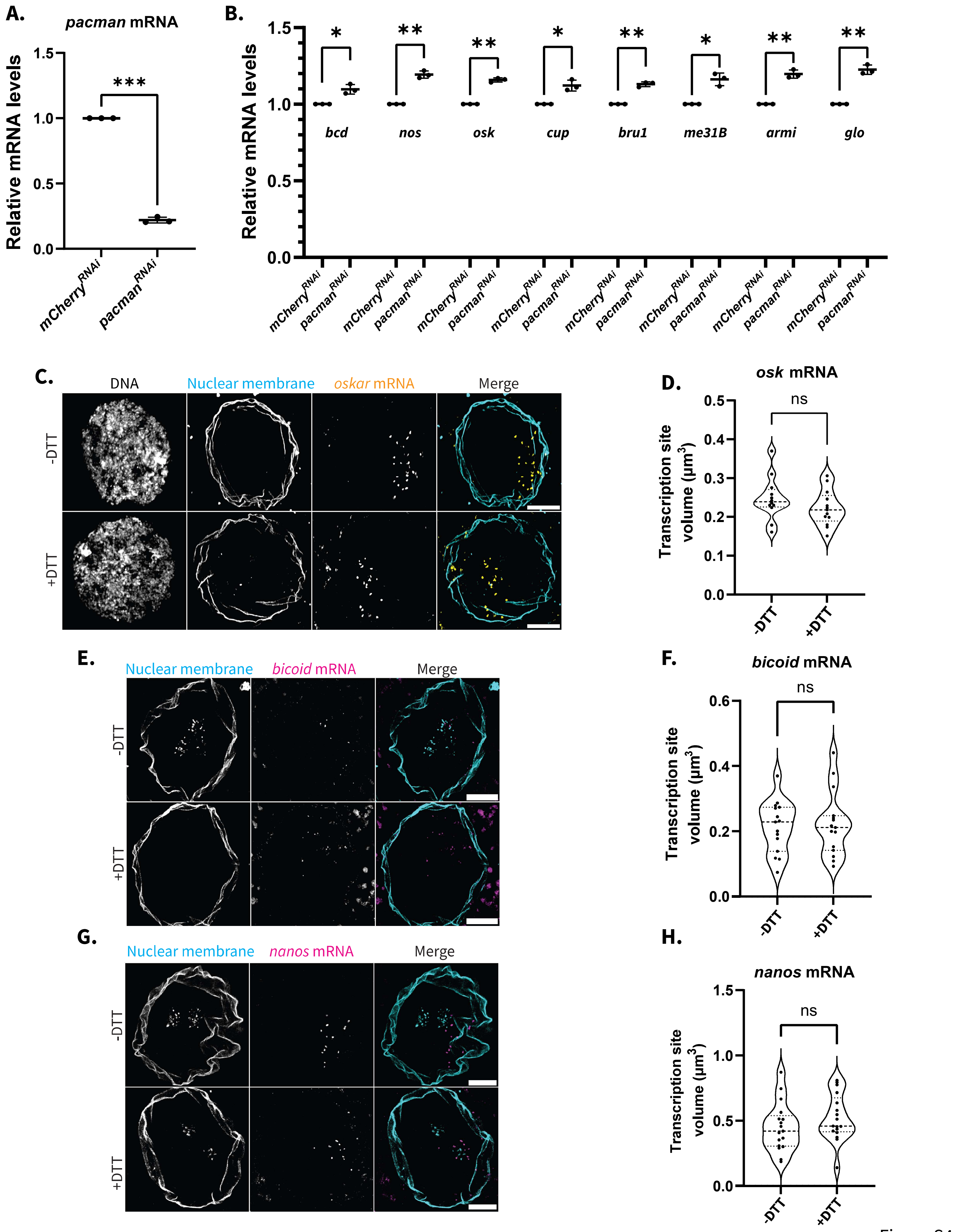
A.
pacman mRNA
Relative mRNA levels
***
mCherry^RNAi
pacman^RNAi
B.
Relative mRNA levels
*
**
**
*
**
*
**
**
bcd
nos
osk
cup
bru1
me31B
armi
glo
C.
DNA
Nuclear membrane
oskar mRNA
Merge
-DTT
+DTT
D.
osk mRNA
Transcription site volume (µm³)
ns
E.
Nuclear membrane
bicoid mRNA
Merge
F.
bicoid mRNA
G.
Nuclear membrane
nanos mRNA
Merge
H.
nanos mRNA

Figure S4

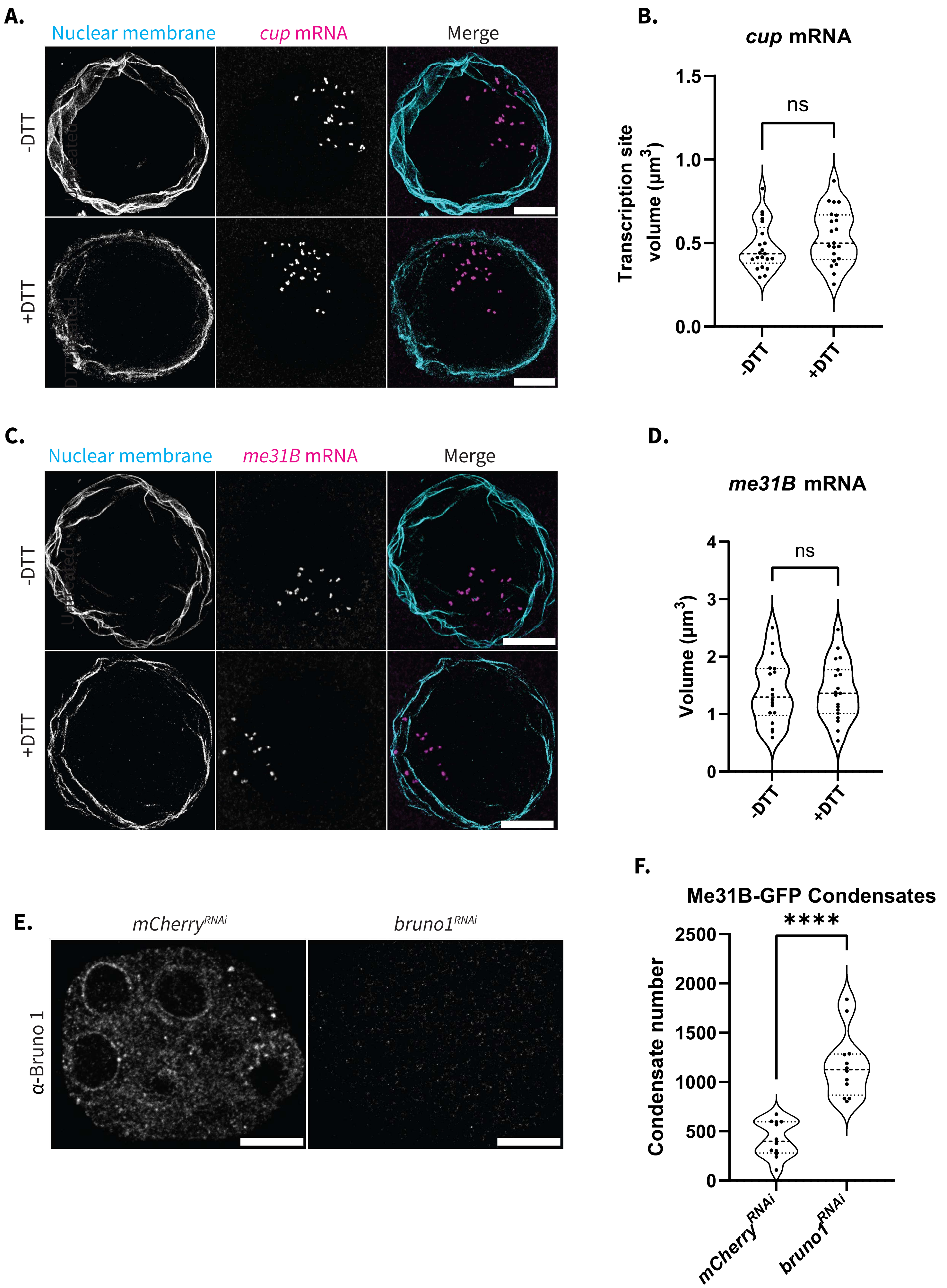
A.
Nuclear membrane
cup mRNA
Merge
-DTT
+DTT
B.
cup mRNA
Transcription site volume (µm3)
ns
1.5
1.0
0.5
0.0
-DTT
+DTT
C.
Nuclear membrane
me31B mRNA
Merge
-DTT
+DTT
D.
me31B mRNA
Volume (µm3)
ns
4
3
2
1
0
-DTT
+DTT
E.
mCherryRNAi
bruno1RNAi
α-Bruno 1
F.
Me31B-GFP Condensates
Condensate number
****
2500
2000
1500
1000
500
0
mCherryRNAi
bruno1RNAi

Figure S5

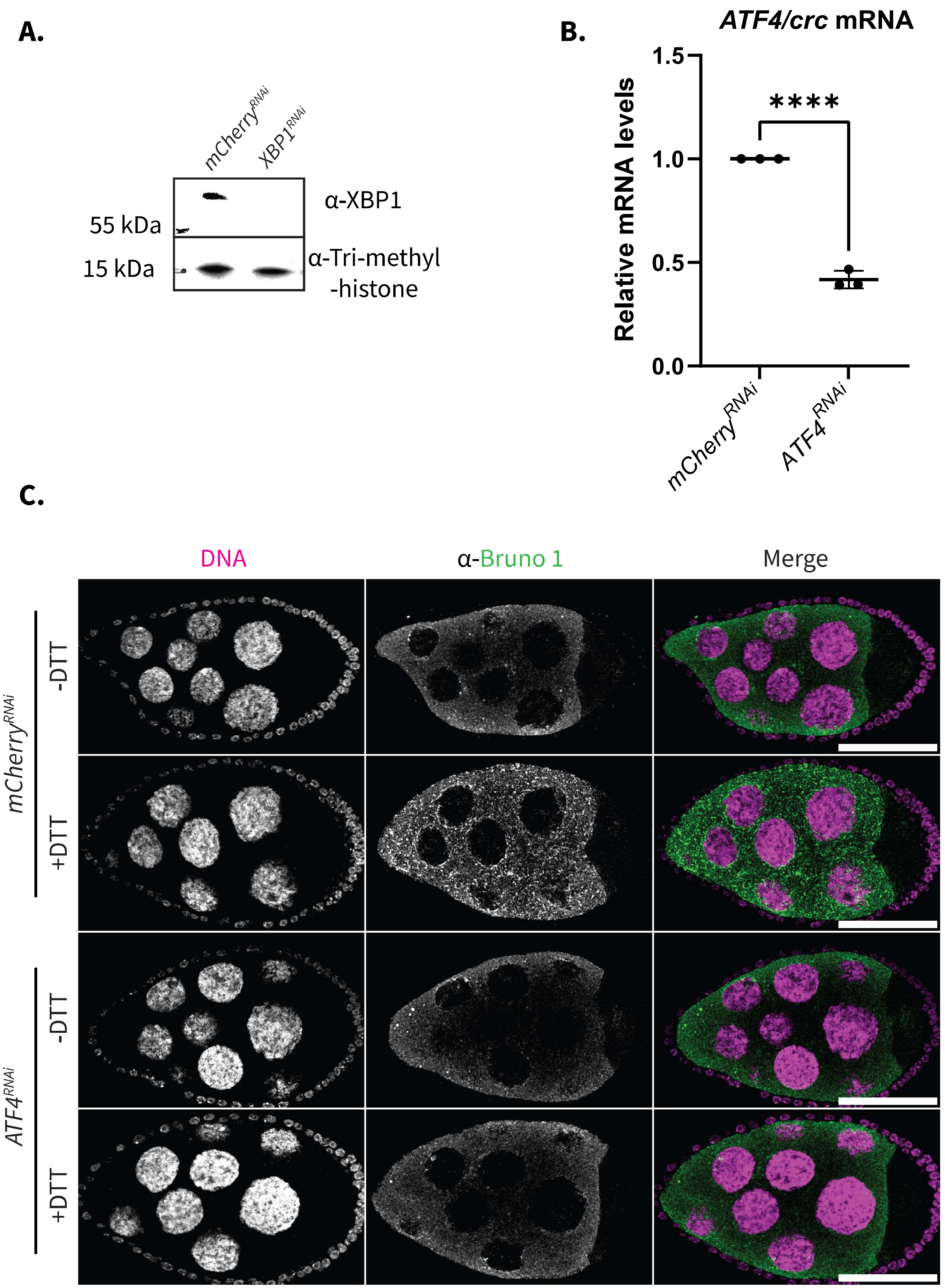
A.
mCherry^RNAi
XBP1^RNAi
55 kDa
15 kDa
α-XBP1
α-Tri-methyl
-histone
B.
ATF4/crc mRNA
Relative mRNA levels
1.5
1.0
0.5
0.0
****
mCherry^RNAi
ATF4^RNAi
C.
DNA
α-Bruno 1
Merge
mCherry^RNAi
ATF4^RNAi
-DTT
+DTT
-DTT
+DTT

Figure S6

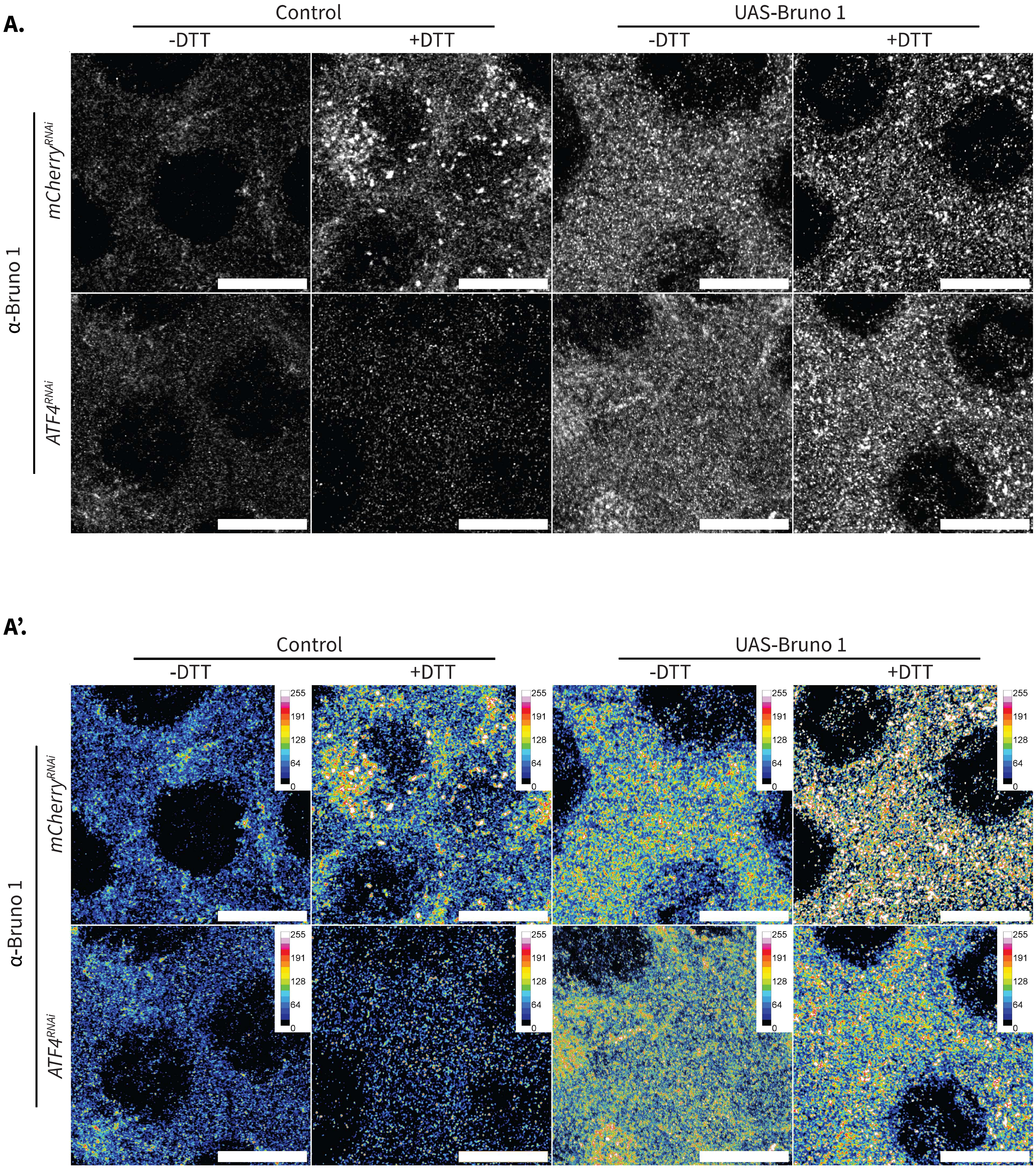
A.
Control
UAS-Bruno 1
-DTT
+DTT
-DTT
+DTT
α-Bruno 1
mCherry^RNAi
ATF4^RNAi
A'.
Control
UAS-Bruno 1
-DTT
+DTT
-DTT
+DTT
α-Bruno 1
mCherry^RNAi
ATF4^RNAi
255
191
128
64
0

Figure S7

**"ATF4-dependent upregulation of Bruno 1 remodels P-bodies to selectively protect mRNAs during ER stress throughout *Drosophila melanogaster* oogenesis"** Milano et al.

---

### SUPPLEMENTAL FIGURES:

**Figure S1:**

**(A)** Visualization of endogenously expressed Me31B-GFP in egg chambers incubated *ex vivo* in Schneider's media alone (-Thapsigargin) or in Schneider's media supplemented with Thapsigargin (+Thapsigargin). Images are XY projections of 5 optical Z slices of 0.3µm. Scale bars are 20µm.

**(B)** Volume quantifications comparing Me31B-GFP condensates in -Thapsigargin and +Thapsigargin incubated egg chambers (n =23).

**(C)** Sphericity quantifications comparing Me31B-GFP condensates in -Thapsigargin and +Thapsigargin incubated egg chambers (n =19).

**(D)** Western blot analysis of Me31B in -DTT and +DTT incubated egg chambers. (Tri-methyl-Histone -- loading control).

**(E)** Visualization of endogenous Cup-YFP condensates in -DTT and +DTT incubated egg chambers. Images are XY projections of 5 optical Z slices of 0.3µm. Scale bars are 20µm.

**(F)** Volume quantifications comparing Cup-YFP condensates in -DTT and +DTT incubated egg chambers (n =20).

**(G)** Sphericity quantifications comparing Cup-YFP condensates in -DTT and +DTT incubated egg chambers (n =20).

**(H)** Visualization of endogenous Tral-RFP condensates in -DTT and +DTT incubated egg chambers. Images are XY projections of 5 optical Z slices of 0.3µm. Scale bars are 20µm.

**(I)** Volume quantifications comparing Tral-RFP condensates in egg chambers incubated for 30 min in -DTT and +DTT (n =20).

**(J)** Sphericity quantifications comparing Tral-RFP condensates in egg chambers incubated for 30 min in -DTT and +DTT (n =20).

For all plots, each data point represents the average value of all P-bodies detected in an image. Significance was assessed using Mann-Whitney statistical tests. Error bars represent standard deviation. **** $P < .0001$.

**Figure S2:**

Covisualization of endogenously expressed Rin-GFP and Tral-RFP in -DTT incubated egg chambers over a 60-minute time course. Images are XY projections of 5 optical Z slices of 0.3µm. Scale bars are 20µm.

**Figure S3:**

**(A-H)** Vantage plots showing the shortest distance from respective mRNA particles to a Me31B-labeled P-body on the X-axis and showing the volume of each mRNA particle on the Y-axis. (n= 3 images, all mRNA particles are represented from each image).

**(A)** *oskar* mRNA.

**(B)** *bicoid* mRNA.

**(C)** *nanos* mRNA.

**(D)** *me31B* mRNA.

**(E)** *cup* mRNA.

**(F)** *bruno1* mRNA.

**(G)** *armi* mRNA.

**(H)** *glorund* mRNA.

**Figure S4:**

**(A)** RT-qPCR quantification of *pacman* mRNAs in *mCherry*$^{RNAi}$ and *pacman*$^{RNAi}$ egg chambers. Significance calculated with Welch's t-test (n = 3).

**(B)** RT-qPCR quantification of 8 mRNAs in *mCherry*$^{RNAi}$ and *pacman*$^{RNAi}$ egg chambers. Significance calculated with Welch's t-test (n = 3).

**(C)** Covisualization of *oskar* mRNA labeled with smFISH probes with DAPI labeled DNA and nuclear membrane stained with wheat agglutinin in -DTT and +DTT incubated egg chambers. Images are XY projections of 15 optical Z slices of 0.3µm. Scale bars are 10µm.

**(D)** Volume quantification of the *oskar* transcription sites in (C) (n=30).

**(E)** Covisualization of *bicoid* mRNA labeled with smFISH probes with the nuclear membrane stained with wheat agglutinin in -DTT and +DTT incubated egg chambers. Images are XY projections of 15 optical Z slices of 0.3µm. Scale bars are 10µm.

**(F)** Volume quantification of the *bicoid* transcription sites in (E) (n=15).

**(G)** Covisualization of *nanos* mRNA labeled with smFISH probes with the nuclear membrane stained with wheat agglutinin in -DTT and +DTT incubated egg chambers. Images are XY projections of 15 optical Z slices of 0.3µm. Scale bars are 10µm.

**(H)** Volume quantification of the *nanos* transcription sites in (G) (n=17).

For all plots, each data point represents the average value of all transcription sites detected in an image. Significance was assessed using Mann-Whitney statistical tests. Error bars represent standard deviation.

**** P < .0001.

**Figure S5:**

**(A)** Covisualization of *cup* mRNA labeled with smFISH probes with the nuclear membrane stained with wheat agglutinin in -DTT and +DTT incubated egg chambers. Images are XY projections of 15 optical Z slices of 0.3μm. Scale bars are 10μm.

**(B)** Volume quantification of the *cup* transcription sites in (A) (n=21).

**(C)** Covisualization of *me31B* mRNA labeled with smFISH probes with the nuclear membrane stained with wheat agglutinin in -DTT and +DTT incubated egg chambers. Images are XY projections of 15 optical Z slices of 0.3μm. Scale bars are 10μm.

**(D)** Volume quantification of the *me31B* transcription sites in (C) (n=18).

**(E)** Visualization of immunolabeled Bruno 1 in $mCherry^{RNAi}$ and $bruno1^{RNAi}$ egg chambers. Images are XY projections of 5 optical Z slices of 0.3μm. Scale bars are 100μm.

**(F)** Quantification of overall Me31B-GFP condensate number in $mCherry^{RNAi}$ and $bruno1^{RNAi}$ egg chambers (n=12).

For all plots, each data point represents the average value of all transcription sites detected in an image. Significance was assessed using Mann-Whitney statistical tests. Error bars represent standard deviation. **** $P < .0001$.

**Figure S6:**

**(A)** Western blot analysis of XBP1 in $mCherry^{RNAi}$ and $XBP1^{RNAi}$ egg chambers. (Tri-methyl-Histone -- loading control).

**(B)** RT-qPCR quantification of *ATF4/crc* mRNA in $mCherry^{RNAi}$ and $ATF4/crc^{RNAi}$ egg chambers. Significance calculated with Welch's t-test (n = 3).

**(C)** Covisualization of immunolabeled Bruno and Dapi labeled DNA in $mCherry^{RNAi}$ and $ATF4/crc^{RNAi}$ egg chambers incubated in -DTT and +DTT. Images are XY projections of 5 optical Z slices of 0.3μm. Scale bars are 100μm. Error bars represent standard deviation. **** $P < .0001$.

**Figure S7:**

**(A)** Visualization of Bruno 1 in control and Bruno 1 overexpression backgrounds (UAS-Bruno 1) in $mCherry^{RNAi}$ and $ATF4/crc^{RNAi}$ egg chambers incubated in -DTT and +DTT. Images are XY projections of 5 optical Z slices of 0.3μm. Scale bars are 20μm.

**(A')** Intensity heat map of **(A)**.
